# Targetable leukemia dependency on noncanonical PI3Kγ signaling

**DOI:** 10.1101/2023.12.15.571909

**Authors:** Qingyu Luo, Evangeline G. Raulston, Miguel A. Prado, Xiaowei Wu, Kira Gritsman, Kezhi Yan, Christopher A. G. Booth, Ran Xu, Peter van Galen, John G. Doench, Shai Shimony, Henry W. Long, Donna S. Neuberg, Joao A. Paulo, Andrew A. Lane

**Affiliations:** Department of Medical Oncology, Dana-Farber Cancer Institute, Harvard Medical School, Boston, MA 02215; Instituto de Investigación Sanitaria del Principado de Asturias (ISPA), Oviedo, Spain; Department of Cell Biology, Harvard Medical School, Boston, MA 02115; Department of Cancer Biology, Dana-Farber Cancer Institute, Boston, MA 02215; Department of Genetics, Blavatnik Institute, Harvard Medical School, Boston, MA 02215; Department of Medical Oncology, Albert Einstein College of Medicine, Bronx, NY 10461; Department of Cell Biology, Albert Einstein College of Medicine, Bronx, NY 10461; Division of Hematology, Brigham and Women’s Hospital, Boston, MA 02115; Genetic Perturbation Platform, Broad Institute of MIT and Harvard, Cambridge, MA 02142; Department of Hematology, Rabin Medical Center, Tel Aviv Faculty of Medicine, Tel Aviv, Israel; Center for Functional Cancer Epigenetics, Dana-Farber Cancer Institute, Boston, MA 02215; Department of Data Science, Dana-Farber Cancer Institute, Boston, MA 02215

## Abstract

Phosphoinositide 3-kinase gamma (PI3Kγ) is implicated as a target to repolarize tumor-associated macrophages and promote anti-tumor immune responses in solid cancers. However, cancer cell-intrinsic roles of PI3Kγ are unclear. Here, by integrating unbiased genome-wide CRISPR interference screening with functional analyses across acute leukemias, we define a selective dependency on the PI3Kγ complex in a high-risk subset that includes myeloid, lymphoid, and dendritic lineages. This dependency is characterized by innate inflammatory signaling and activation of phosphoinositide 3-kinase regulatory subunit 5 (*PIK3R5*), which encodes a regulatory subunit of PI3Kγ and stabilizes the active enzymatic complex. Mechanistically, we identify p21 (RAC1) activated kinase 1 (PAK1) as a noncanonical substrate of PI3Kγ that mediates this cell-intrinsic dependency independently of Akt kinase. PI3Kγ inhibition dephosphorylates PAK1, activates a transcriptional network of NFκB-related tumor suppressor genes, and impairs mitochondrial oxidative phosphorylation. We find that treatment with the selective PI3Kγ inhibitor eganelisib is effective in leukemias with activated *PIK3R5*, either at baseline or by exogenous inflammatory stimulation. Notably, the combination of eganelisib and cytarabine prolongs survival over either agent alone, even in patient-derived leukemia xenografts with low baseline PIK3R5 expression, as residual leukemia cells after cytarabine treatment have elevated G protein-coupled purinergic receptor activity and PAK1 phosphorylation. Taken together, our study reveals a targetable dependency on PI3Kγ/PAK1 signaling that is amenable to near-term evaluation in patients with acute leukemia.

## Introduction

Despite recent progress with development of targeted therapies for some genetic subsets of acute leukemias^1,2^, many disease subtypes lack mechanistically targeted treatment, and most patients have poor long-term outcomes. Acute myeloid leukemia (AML) and acute lymphocytic leukemia (ALL) are the most common types of acute leukemia in adults and children, respectively. Blastic plasmacytoid dendritic cell neoplasm (BPDCN) is a rare and aggressive hematologic malignancy that originates from the plasmacytoid dendritic cell (pDC) lineage and shares clinical and pathologic features with both AML and ALL^3,4^. Therapy directed at cell surface-expressed CD123 is approved in BPDCN and is being tested in other acute leukemias^5,6^, but it does not directly target cell-intrinsic oncogenic pathways. We hypothesized that overlapping features between BPDCN and other acute leukemias may indicate a functionally related subset of hematologic malignancies that have shared therapeutic vulnerabilities.

The mammalian phosphoinositide 3-kinase (PI3K) family contains 8 isoforms that can be divided into several classes. Class I PI3Ks, divided into class IA and IB, generate 3-phosphoinositide lipids to activate signal transduction pathways, while class II and class III PI3Ks are regulators of membrane traffic along the endocytic route^7^. Class IA PI3Ks include three catalytic subunits α, β, and δ (encoded by *PIK3CA*, *PIK3CB*, and *PIK3CD*) with regulatory subunits p85α, p55α, p50α, p85β, and p55γ (encoded by *PIK3R1*, *PIK3R2*, and *PIK3R3*)^7^. Class IB PI3K has only one catalytic subunit p110γ (encoded by *PIK3CG*) and 2 regulatory subunits p101 and p84 (encoded by *PIK3R5* and *PIK3R6*), and is activated by G protein-coupled receptors (GPCRs) via heterotrimeric and small G proteins^8^. Several cancers harbor activated class IA PI3Ks, and numerous pathway inhibitors are approved or in development^9,10^. In contrast, Class IB PI3K components (i.e., the enzymatic p110γ and regulatory subunit(s)) have received less attention and therapeutic focus has been limited to reprograming macrophages with PI3Kγ inhibitors for solid tumor immunotherapy^11–14^. The role of PI3Kγ as a cell-intrinsic cancer driver remains unclear.

Developments in functional genomics, including updated CRISPR technologies, facilitate unbiased screening and integration of data across cancer types^15^. An RNA interference screen using shRNA was previously performed in BPDCN cell lines, but it involved a relatively small pre-selected set of genes, and the hits were mostly known pDC lineage dependencies^16^. Here, through genome-wide CRISPR interference (CRISPRi) screening followed by integrated analyses, we define a dependency on noncanonical PI3Kγ signaling that is shared by the majority of BPDCNs and a proportion of other acute leukemias (AMLs and ALLs). We find that PI3Kγ signaling is activated by the elevation of *PIK3R5* and drives malignant phenotypes through phosphorylation of p21 (RAC1) activated kinase 1 (PAK1), sustaining mitochondrial oxidative phosphorylation (OXPHOS), and inhibiting NFκB-related tumor suppressor genes. This PIK3R5-driven signaling pathway is sensitive to PI3Kγ inhibition, providing a novel therapeutic strategy for these aggressive leukemias.

## Results

### Integrated CRISPRi and bioinformatic screening reveals leukemia intrinsic dependency on the PI3Kγ complex

With the hypothesis that a shared targetable dependency exists between BPDCN and other aggressive leukemias, we utilized CAL-1, a BPDCN cell line, for a genome-wide CRISPRi dependency screen (**Fig. 1a**). We tested the Dolcetto guide RNA sets A and B, which include 114,061 guides targeting ∼19,000 genes (average 6 guides/gene)^15^. Samples were collected at day 0, 14, and 21 post-puromycin selection (**Fig. 1a**). As expected for positive controls, the top dependencies at day 14 and 21 belong to ribosomal protein and/or Cancer Dependency Map (DepMap) common essential genes, which are required for survival of most human cells^17^ (**Fig. 1b**). Among 1335 significant CAL-1 dependencies, we defined 327 genes as core dependencies with a strict threshold of *P* < 0.001 and fold change < 0.5 (2-fold depletion) comparing day 21 to day 0 (**Fig. 1b**). No gene in the day 21 dependencies overlapped with the nonessential negative control gene list from DepMap, while 143 out of the 327 core dependencies were also DepMap common essentials, validating the screen output (**Fig. 1c**). Therefore, the 184 genes not overlapping with the common essential list represented potential specific CAL-1/BPDCN dependencies (**Fig. 1c**). Unlike the 143 common essential genes, the 184 specific dependency genes were highly enriched for pathways implicated in AML, supporting the relatedness of BPDCN and AML (**Fig. 1c,d**).

**Figure 1.**
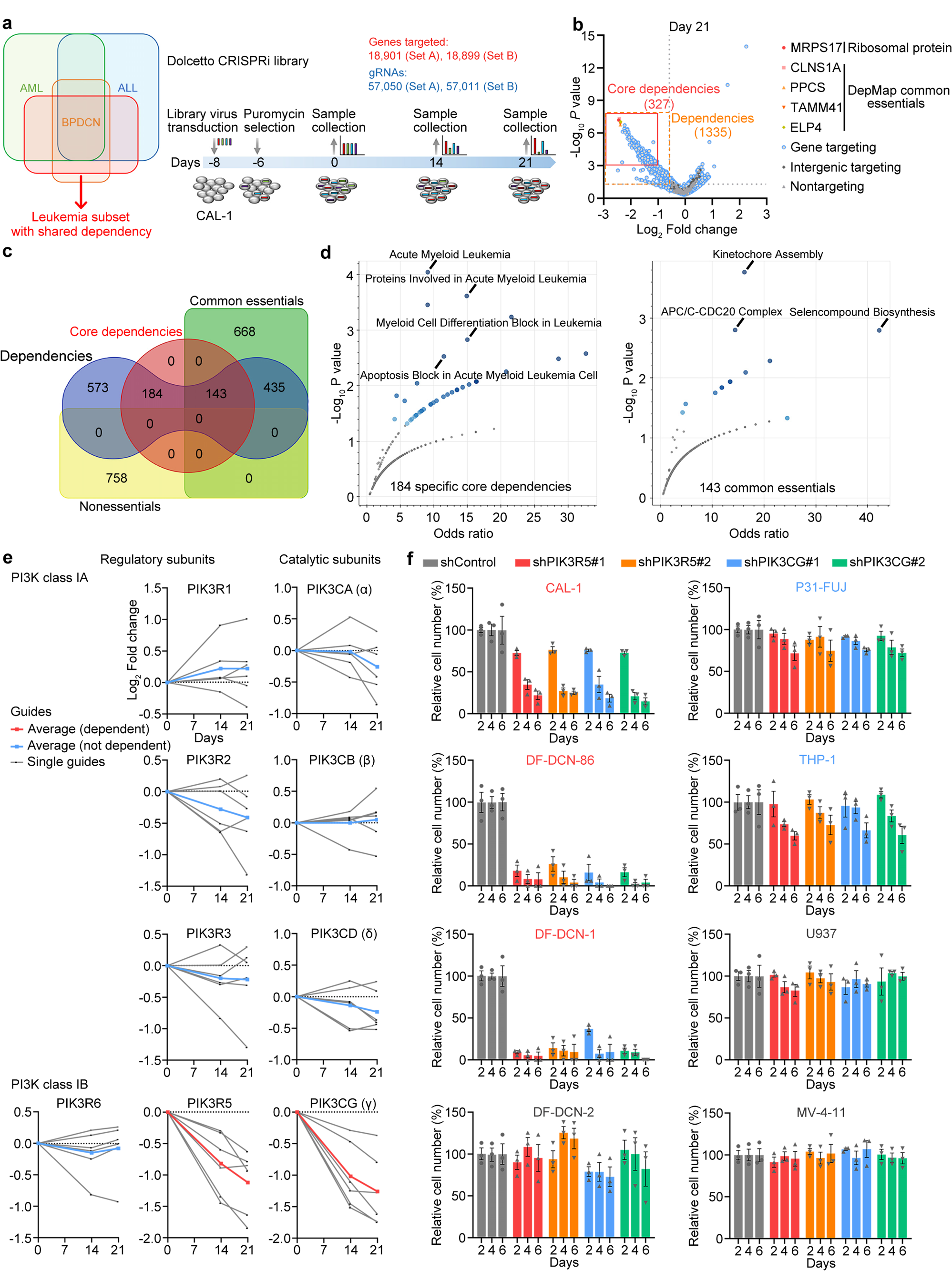
Genome-wide CRISPRi screening identifies a selective dependency of BPDCN on PI3Kγ. **a,** Schematics of the concept and experimental design to identify a targetable dependency shared by aggressive acute leukemias including BPDCN and other pathologic subtypes. **b,** Volcano plot showing the depletion of all genes at day 21. Orange square indicates dependency genes (*P* < 0.05 and depletion fold >1.5), and red square indicates core dependency genes (*P* < 0.001 and depletion fold >2). **c,** Overlapping gene numbers between the day 21 dependencies and common essential as well as nonessential genes from DepMap. **d,** Significantly enriched pathways with the Elsevier Pathway Collection via Enrichr analyses of the 184 specific core dependencies or the 143 common essentials. **e,** Depletion of guides targeting PI3K subunits in the screen. **f,** Relative cell numbers of BPDCN and AML cell lines after depletion of PIK3R5 or PIK3CG at 2, 4, and 6 days as compared to control cells. Strongly sensitive cell lines are labelled in red, moderately sensitive cell lines are labelled in blue, and insensitive cell lines are labelled in grey. Data are means ± S.E.M. from 3 biological replicates.

We next asked if a dependency was shared by BPDCN and only a subset of AML with similar biology, reasoning that such a therapeutic target may not have been identified in prior studies focused on AML more generally. We used the following criteria to narrow the list of 184 day 21 specific dependencies: 1. Not a dependency in all or most AMLs in DepMap (no more than 16 out of 23 AML cell lines); 2. Highly expressed in BPDCN compared to normal pDCs according to our previous RNA-seq data^18^; 3. Highly expressed in BPDCN compared to most AMLs according to a side-by-side expression analysis of >200 blood cancer patient-derived xenograft (PDX) models^19^. Following these criteria, only one gene, *PIK3R5*, was a hit (**Extended Data Fig. 1a-c**). Of note, a previously identified pDC lineage dependency gene *TCF4*^16^ was also in our dependency gene list and showed higher expression in BPDCN than in other leukemias but showed no difference in its expression between BPDCN and normal pDCs (**Extended Data Fig. 1b,c**). *PIK3R5* encodes the p101 regulatory subunit (hereafter, referred to as PIK3R5) of the PI3Kγ complex. *PIK3CG*, which encodes the PI3Kγ complex catalytic subunit p110γ (hereafter, PIK3CG) and was a strong hit in CAL-1 (**Fig. 1e**), did not pass our second criterion as it is a weak-moderate dependency in most AML cell lines (**Extended Data Fig. 1a**). In contrast, none of the other PI3K complex components was a dependency gene from our screen, suggesting a selective dependency of BPDCN and related leukemias on the class IB PI3Kγ complex (**Fig. 1e**). To validate PI3Kγ complex dependency from independent experiments, we analyzed *PIK3R5* and *PIK3CG* across all cancer cell lines in DepMap. Albeit with moderate scores, the PI3Kγ complex was selectively required by some blood cancers but not any solid tumors, and the same cells were dependent on both *PIK3R5* and *PIK3CG*, supporting their relationship in dependent cells (**Extended Data Fig. 2a**).

To perform additional functional validation, we first needed to overcome the lack of widely available BPDCN cell lines, which has in part been related to the difficulty in establishing cultures from patient material. We created a leukemia adaptive medium (LAM), modified from a previously established organoid culture regimen^20^, which enabled BPDCN primary patient samples to maintain viability for at least two weeks ex vivo. By gradually introducing standard leukemia culture medium (see Methods), we successfully established a new BPDCN cell line named DF-DCN-86 (**Extended Data Fig. 2b**). Since the patient sample for the establishment of DF-DCN-86 was a BPDCN skin biopsy, we xenografted DF-DCN-86 cells intradermally into NSG mice and observed the formation of skin tumors that resemble the original patient pathology (**Extended Data Fig. 2c,d**). Using a similar methodology, we established two additional BPDCN cell lines, DF-DCN-1 and DF-DCN-2, from independent patient bone marrow samples.

Three of four BPDCN cell lines exhibited elevated PIK3R5 (mRNA and protein) and PIK3CG (protein only) compared to AML cell lines, consistent with our in silico analysis (**Extended Data Fig. 2e,f**). All leukemia cell lines with high PIK3R5/PIK3CG were strikingly sensitive to PIK3R5/PIK3CG knockdown, while cells with moderate or low PIK3R5/PIK3CG were moderately or not sensitive, respectively (**Fig. 1f and Extended Data Fig. 3a**). We also tested several PI3K inhibitors (eganelisib, a selective PI3Kγ inhibitor; duvelisib, a PI3Kδ and PI3Kγ dual inhibitor; and LY294002, a PI3Kα, PI3Kδ, and PI3Kβ inhibitor). Again, the results indicated strong sensitivity of the three leukemia cell lines with the highest PIK3R5/PIK3CG expression to the selective PI3Kγ inhibitor eganelisib (**Extended Data Fig. 3b**). Modest sensitivity was observed for duvelisib, while no correlation was observed between PIK3R5/PIK3CG expression and sensitivity to LY294002 (**Extended Data Fig. 3b**). These results suggested that PIK3R5/PIK3CG expression is a specific indicator of PI3Kγ dependency.

### PIK3R5 expression is correlated with an innate immune response signature (IIRS) and can be activated by innate inflammatory signaling

To further define PI3Kγ complex dependency, we first tested the vulnerability of AML, ALL, and BPDCN PDX models^19^ to PI3Kγ inhibition. All leukemias with elevated PIK3R5 showed sensitivity to eganelisib, while those with low PIK3R5 expression did not (**Fig. 2a and Extended Data Fig. 4a,b**). Supporting induction of apoptosis as the mechanism of action, eganelisib treatment activated caspase-3/7 in sensitive PIK3R5-high leukemias but not in PIK3R5-low cases (**Extended Data Fig. 4c**). In addition, PI3Kγ inhibition induced markers of terminal myeloid differentiation (CD11b and CD14) in AMLs with high PIK3R5 (**Extended Data Fig. 4d**). Taken together, these results indicated a leukemia intrinsic dependency on the PI3Kγ complex shared by various pathological subtypes that can be predicted by PIK3R5 mRNA expression level.

**Figure 2.**
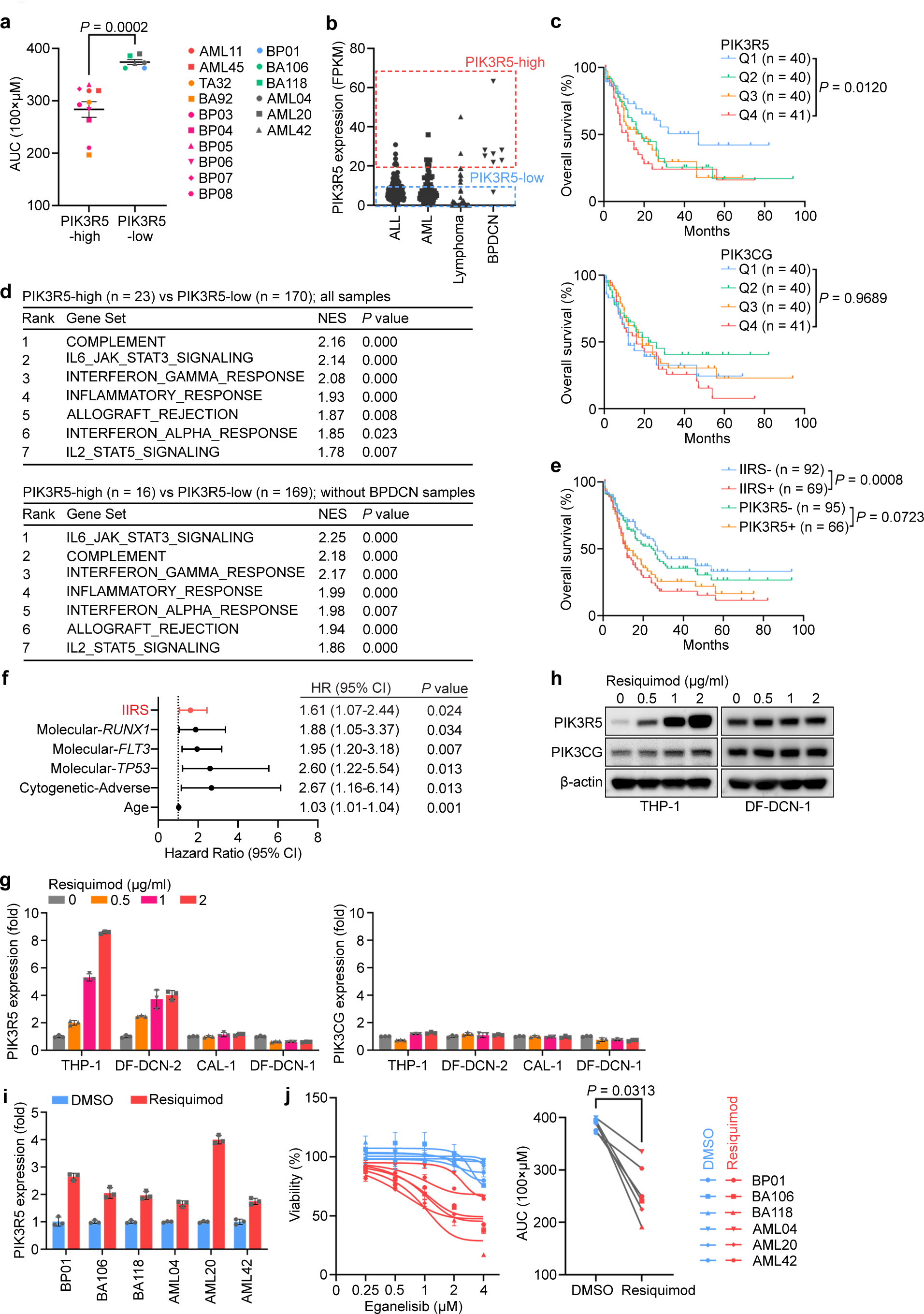
Selective dependency on PI3Kγ subunits is shared by a leukemia subset that is defined by innate inflammatory signaling activated *PIK3R5*. **a,** Area under the curve (AUC) of leukemia PDX samples after ex vivo treatment with increasing doses of eganelisib (0-4 μM) for 72 hours. Data are means ± S.E.M. with Mann-Whitney test. **b,** PIK3R5 mRNA expression level of various blood cancer PDXs from RNA-seq. **c,** Overall survival of AML patients from TCGA. Log-rank test (Q4 vs Q1). **d,** Top enriched gene sets from GESA comparing high PIK3R5 (PIK3R5-high; FPKM>20) to low PIK3R5 (PIK3R5-low; FPKM<10) expression groups. **e,** Overall survival of AML patients from TCGA. Log-rank test. **f,** Hazard ratios from multivariable analysis of factors independently associated with AML prognosis. IIRS: high vs low; *RUNX1*, *FLT3*, and *TP53*: mutated vs nonmutated; Cytogenetic-adverse vs favorable; Age: as continuous variable. **g,** RT-qPCR of PIK3R5 and PIK3CG mRNA in leukemia cell lines after resiquimod treatment for 48 hours. Data are means ± S.D. from 3 technical replicates. **h,** Western blotting for PIK3R5 and PIK3CG in leukemia cell lines after resiquimod treatment for 48 hours. **i,** RT-qPCR of PIK3R5 mRNA in leukemia PDXs after ex vivo resiquimod treatment (1 μg/ml) for 48 hours. Data are means ± S.D. from 3 technical replicates. **j,** Inhibition curve and AUC of leukemia PDXs after ex vivo treatment with increasing doses of eganelisib (0-4 μM) for 72 hours, with or without resiquimod (1 μg/ml). Data are means ± S.D. from 3 technical replicates (left); Mann-Whitney test (right).

Next, we asked if there was a common genetic feature among leukemias with PI3Kγ dependency. Among the patient-derived models (PDXs and cell lines), there was no evidence of a recurrent mutation(s) clearly associated with complex dependency or sensitivity to eganelisib. Therefore, we hypothesized that this PI3Kγ dependency was predominantly determined by transcriptional activation of *PIK3R5*. To support this hypothesis, we divided leukemia PDXs into three groups based on PIK3R5 expression: PIK3R5-high, PIK3R5-mid, and PIK3R5-low. This defined a leukemia subset with high PIK3R5 mRNA that included the majority (7 out of 8) of BPDCNs plus ∼10% of other acute leukemias (**Fig. 2b**). Supporting potential clinical relevance, analysis of AML patient survival data from The Cancer Genome Atlas (TCGA) showed that the highest PIK3R5 patient quartile exhibited significantly worse prognoses than those with the lowest expression (**Fig. 2c**). On the other hand, expression of PIK3CG showed no association with AML patient survival (**Fig. 2c**), further supporting our model that PIK3R5 is the predictive marker of this leukemia subset.

To further elucidate characteristics of leukemias with elevated PIK3R5, we performed gene set enrichment analysis (GSEA) comparing PIK3R5-high to PIK3R5-low cases. Intriguingly, all the top enriched gene sets were related to innate immune response pathways (**Fig. 2d**). The same conclusion was obtained after removing BPDCN cases from the analysis, excluding the potential bias that might be caused by the dendritic cell origin of BPDCN (**Fig. 2d**). We next defined an innate immune response signature (IIRS) of 32 genes that were part of the GSEA core enrichment in at least three out of the seven top gene sets (**Fig. 2d and Extended Data Fig. 4e**). Analysis of TCGA AML patients showed that the IIRS score highly correlated with PIK3R5 mRNA expression (**Extended Data Fig. 4f**) and was more significantly associated with overall survival than the PIK3R5 single gene score (**Fig. 2e**). To further determine the contribution of the IIRS score in predicting prognosis, we performed multivariable analysis of clinical and pathologic factors in the TCGA cohort. We found that the IIRS score was significantly associated with poor prognosis, independent of age, cytogenetics, or somatic mutations (**Fig. 2f**). Again, no recurrent mutation was specifically enriched in IIRS positive group (**Extended Data Fig. 4g**). In support of the IIRS defining a potentially functional leukemia subset associated with inflammatory characteristics, French-American-British M4 and M5 (myelo-monocytic and monocytic) pathologic subtypes of AML were enriched in TCGA cases with high IIRS (52.2% vs 19.7%, *P* < 0.0001). Together, these results show that PIK3R5 expression and the IIRS score predict PI3Kγ dependency and may define a inflammation-associated subset of leukemia.

To further define the relationship between innate inflammatory signaling and PIK3R5 expression, we treated leukemia cell lines, two each with low or high PIK3R5, with resiquimod. Resiquimod is an agonist of Toll-like receptors (TLRs), pattern recognition receptors that initiate innate immune responses by sensing microbial stimuli and are also tonically activated in some hematologic malignancies^21–24^. Resiquimod treatment strikingly increased PIK3R5 mRNA and protein levels in leukemia cells with baseline low PIK3R5 expression (THP1 and DF-DCN-2; **Fig. 2g**). In contrast, PIK3R5 was not further activated in leukemia cells with baseline elevated PIK3R5 (CAL-1 and DF-DCN-1; **Fig. 2g**). PIK3CG mRNA levels were unchanged after resiquimod treatment in all cell lines, but PIK3CG protein levels increased in parallel with PIK3R5 (**Fig. 2h**). Finally, we determined whether resiquimod-induced PIK3R5 elevation could induce sensitivity to PI3Kγ inhibition. We found that resiquimod treatment increased PIK3R5 expression and sensitivity to eganelisib in all six leukemia PDXs with baseline low PIK3R5 (**Fig. 2i,j**). Together, these results support a model that activation of innate inflammatory signaling contributes to elevated PIK3R5 expression and to dependency of those leukemia cells on the PI3Kγ complex.

### PU.1 activates PIK3R5 to promote a self-stabilizing PI3Kγ complex

We next asked what is the upstream transcriptional activator that promotes elevated PIK3R5 expression and dependency on the PI3Kγ complex. We measured accessible chromatin by assay for transposase accessible chromatin with high-throughput sequencing (ATAC-seq) in eight AML, ALL and BPDCN PDXs. Regardless of leukemia histology, the chromatin regions around the *PIK3R5* transcription start site (TSS) were more accessible in cells with high PIK3R5 than those with low PIK3R5 (**Fig. 3a**). As a negative control, no difference was observed in the *PIK3CG* locus between PIK3R5 high and low groups (**Fig. 3a**). Then, we analyzed chromatin immunoprecipitation followed by sequencing (ChIP-seq) data from the ENCODE database^25^ and identified 34 transcriptional regulators that interact with the *PIK3R5* promoter near its TSS (**Extended Data Fig. 5a**). Among the genes encoding these 34 transcriptional regulators, *SPI1* and *ASH2L* were also core dependency genes from our genome wide CRISPRi screen (**Fig. 1b** and **Extended Data Fig. 5a**). Thus, we hypothesized that one of these transcriptional regulators may control activation of *PIK3R5*.

**Figure 3.**
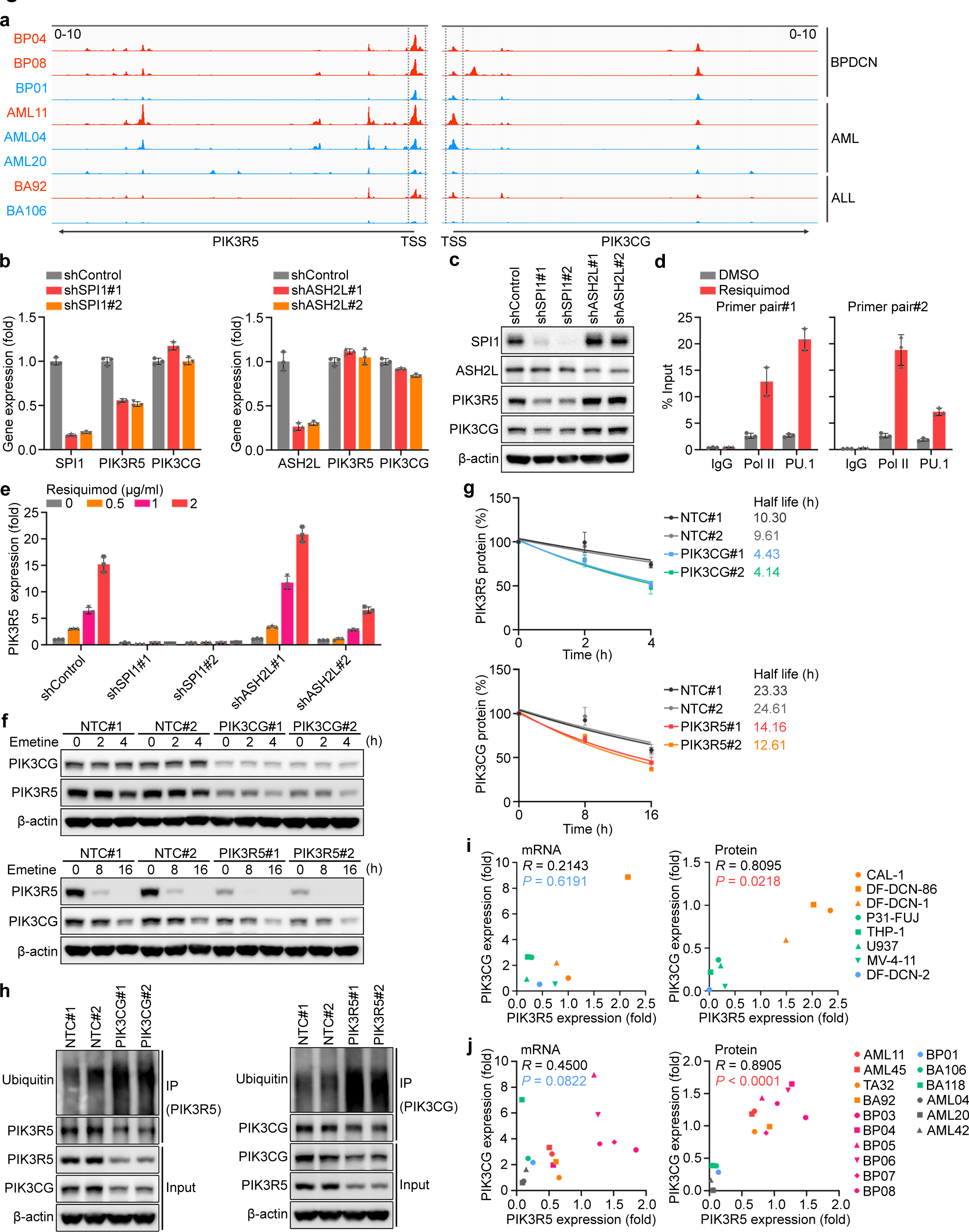
*PIK3R5* is transcriptionally activated by PU.1 and protects the PI3Kγ complex from UPS-mediated degradation. **a,** ATAC-seq peaks at *PIK3R5* and *PIK3CG* genomic regions. **b,c,** RT-qPCR (b) and western blotting (c) analyses of PIK3R5 and PIK3CG expressions in CAL-1 cells with nontargeting (shControl), SPI1 targeting (shSHP1#1, shSPI1#2), or ASH2L targeting (shASH2L#1, shASH2L#2) shRNAs. Data are means ± S.D. from 3 technical replicates. **d,** ChIP detecting the enrichment of RNA polymerase II (Pol II) and PU.1 on *PIK3R5* promoter regions in THP-1 cells treated with vehicle (DMSO) or resiquimod (1 μg/ml) for 48 hours. Data are means ± S.D. from 3 technical replicates. **e,** RT-qPCR of PIK3R5 mRNA in THP-1 cells expressing nontargeting, SPI1 targeting, or ASH2L targeting shRNAs after resiquimod treatment (1 μg/ml) for 48 hours. **f,g,** Western blotting (f) and quantitation (g) for PIK3R5 and PIK3CG protein levels in control (NTC#1, NTC#2), PIK3CG depleted (PIK3CG#1, PIK3CG#2), or PIK3R5 depleted (PIK3R5#1, PIK3R5#2) CAL-1 cells treated with the protein synthesis inhibitor emetine (50 μg/ml) for the indicated times. Data are means ± S.E.M. from 3 biological replicates. **h,** Western blotting for PIK3R5 and PIK3CG ubiquitination in control, PIK3CG depleted, or PIK3R5 depleted CAL-1 cells. **i,j,** Correlation analyses of PIK3R5 and PIK3CG mRNA or protein expressions in leukemia cell lines (i) or PDXs (j). Spearman correlation coefficients are shown.

Therefore, we depleted SPI1 or ASH2L and measured expression of PIK3R5. Knockdown of SPI1, but not ASH2L, substantially reduced the level of PIK3R5 mRNA and protein (**Fig. 3b,c**). *SPI1* encodes the transcription factor PU.1, which has important roles in myeloid and lymphoid differentiation^26,27^, as well as hematopoietic stem cell maintenance^28,29^. In some leukemias, targeting PU.1 suppresses cell growth and triggers apoptosis^30,31^, which is supported by DepMap data that *SPI1* is a selective dependency in some myeloid and lymphoid cells (**Extended Data Fig. 5b**). Additionally, resiquimod induces a PU.1-mediated gene expression program in neutrophils^32^. We next performed ChIP assays and observed that the association between PU.1 and the *PIK3R5* promoter was increased upon resiquimod treatment (**Fig. 3d**). In addition, depletion of SPI1 abolished the effect of resiquimod on *PIK3R5* activation (**Fig. 3e**). These results indicated that PU.1 is a critical transcription factor mediating *PIK3R5* activation in the PI3Kγ-dependent leukemia subset.

Our results showed that transcriptional activation of *PIK3R5* but not *PIK3CG* contributes to the intrinsic dependency of leukemias on the PI3Kγ complex. We noted that the protein level of either PIK3R5 or PIK3CG was reduced upon depletion of the other gene (**Extended Data Fig. 3a**). Additionally, the protein level of PIK3CG increased following the activation of *PIK3R5* despite the PIK3CG mRNA level remaining unchanged (**Fig. 2g,h**). Furthermore, depletion of PIK3R5 or PIK3CG did not affect the other’s mRNA expression (**Extended Data Fig. 5c**). We therefore hypothesized that the PIK3R5 and PIK3CG proteins may protect each other from turnover, increasing their half-life in cells. To test this possibility, we treated CAL-1 cells with the protein synthesis inhibitor emetine and monitored the stability of PIK3R5 and PIK3CG proteins. We observed that depletion of PIK3CG strikingly reduced the stability of PIK3R5 protein, and vice versa (**Fig. 3f,g**). Since the ubiquitin-proteasome system (UPS) mediates degradation of most cellular proteins^33,34^, we performed ubiquitination assays and found that depletion of PIK3R5 or PIK3CG enhances the ubiquitination of the other (**Fig. 3h**). This mechanism of shared protein protection from UPS-mediated degradation was further supported by observing a significant correlation between PIK3R5 and PIK3CG protein levels, but not mRNA levels, in leukemia cell lines (**Fig. 3i and Extended Data Fig. 2e,f**) and PDXs (**Fig. 3j and Extended Data Fig. 4a, 5d,e**).

### PI3Kγ inhibition suppresses OXPHOS and activates an NFκB-related transcriptional network

To study the downstream events underlying leukemia cell-intrinsic PI3Kγ dependency, we performed integrated RNA-seq, proteomic, and phosphoproteomic analyses using CAL-1 cells treated with eganelisib or depleted of PIK3R5 using CRISPRi (PIK3R5i). With inhibition or depletion, GSEA from cell transcriptomes indicated that the most significantly downregulated molecular event after targeting PI3Kγ was related to OXPHOS (**Fig. 4a**). This provides additional rationale to selectively target mitochondria in leukemia cells by inhibiting PI3Kγ, especially in AML, which is typically deficient in glycolysis and dependent on OXPHOS^35^. We performed metabolic phenotyping (Seahorse) assays to validate the effects of PI3Kγ inhibition on cellular energy metabolism. Treatment with eganelisib, but not the class IA PI3K inhibitor LY294002, suppressed OXPHOS in three different BPDCN cell lines (**Fig. 4b**). Suppression of OXPHOS was also observed after CRISPRi-mediated PIK3R5/PIK3CG knockdown, confirming the on-target effect of eganelisib on PI3Kγ (**Fig. 4c**). In contrast, glycolysis was not affected by eganelisib treatment or PIK3R5/PIK3CG knockdown, indicating the effect of PI3Kγ inhibition on energy metabolism is mainly related to suppression of OXPHOS (**Extended Data Fig. 6a,b**).

**Figure 4.**
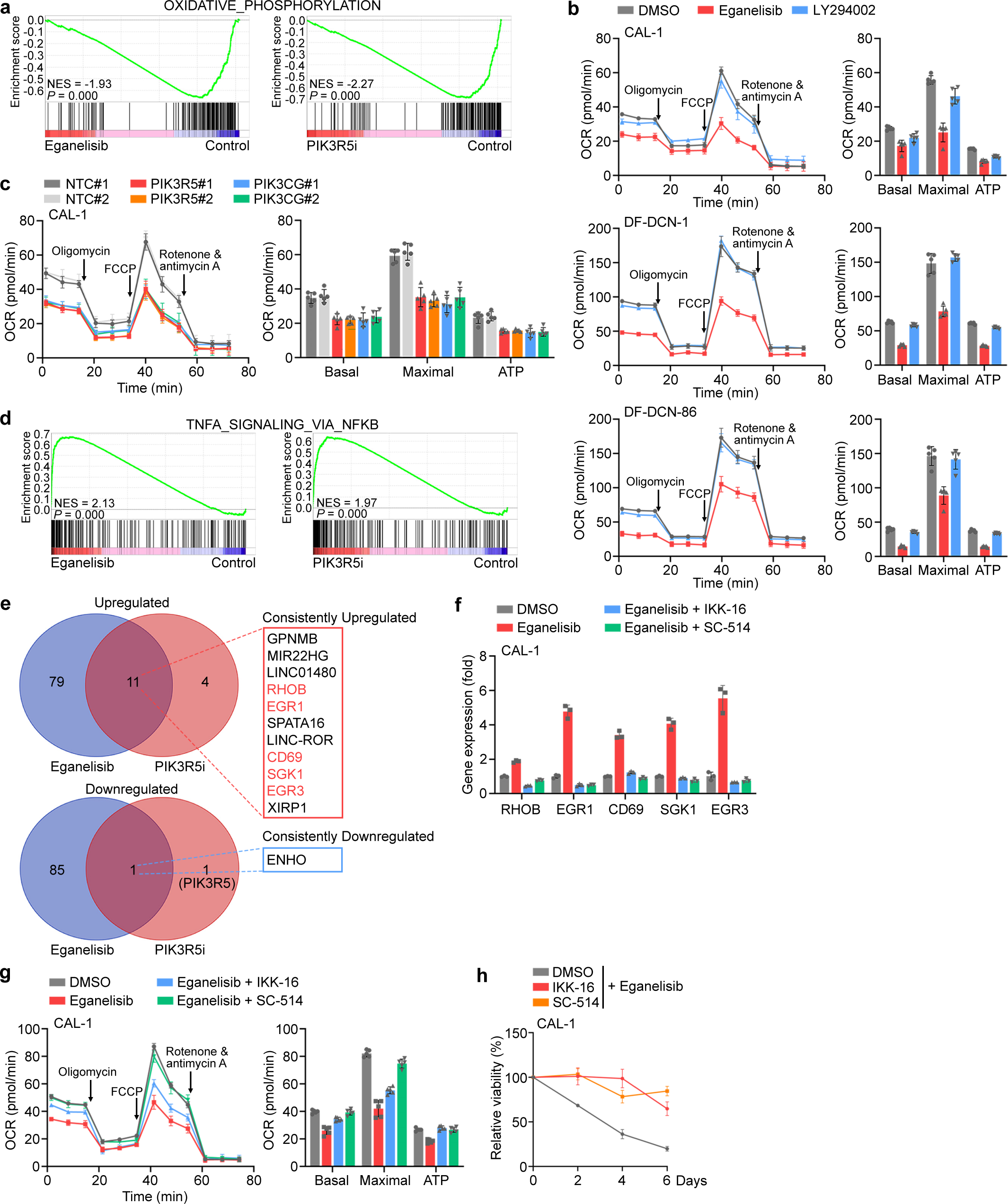
PI3Kγ inhibition activates an NFkB-related transcriptional network and suppresses OXPHOS in leukemia cells. **a,** GSEA indicating OXPHOS to be the most suppressed gene set upon eganelisib treatment (1 μM for 48 hours) or PIK3R5 depletion (PIK3R5i) in CAL-1 cells. **b,** Seahorse assays detecting mitochondrial respiration in the indicated leukemia cell lines upon treatment with eganelisib or LY294002 (each 1 μM for 48 hours). Data are means ± S.D. from 5 technical replicates. **c,** Seahorse assays detecting mitochondrial respiration in control, PIK3R5 depleted, or PIK3CG depleted CAL-1 cells. Data are means ± S.D. from 5 technical replicates. **d,** GSEA results indicating TNFα signaling via NFkB to be the most activated gene set upon eganelisib treatment or PIK3R5 depletion in CAL-1 cells. **e,** Significantly upregulated or downregulated genes in CAL-1 cells treated with eganelisib or depleted of PIK3R5. Five genes belonging to the TNFα signaling via NFkB gene set are labelled in red. **f,** RT-qPCR detection of gene expression in CAL-1 cells treated with eganelisib (1 μM), with or without NFkB inhibitors IKK-16 (0.5 μM) or SC-514 (20 μM), for 2 days. Data are means ± S.D. from 3 technical replicates. **g,** Seahorse assays detecting mitochondrial respiration in CAL-1 cells treated with eganelisib (1 μM), with or without NFkB inhibitors IKK-16 (0.5 μM) or SC-514 (20 μM), for 2 days. Data are means ± S.D. from 5 technical replicates. **h,** Relative CAL-1 viability after treatment with eganelisib (1 μM), with or without NFkB inhibitors IKK-16 (0.5 μM) or SC-514 (20 μM), for 2, 4, and 6 days. Data are means ± S.E.M. from 3 biological replicates.

The most significantly upregulated gene set upon PI3Kγ inhibition was related to TNFα signaling-mediated NFκB activity (**Fig. 4d**). Five of eleven consistently elevated genes (*RHOB*, *EGR1*, *CD69*, *SGK1*, and *EGR3*) after eganelisib treatment and PIK3R5i are within the core enrichment group of this gene set (**Fig. 4e**), and several have tumor suppressive roles in various cancers^36–38^. Thus, we hypothesized that one or more of these core enriched genes contributed to the vulnerability of the PIK3R5-related leukemia subset to PI3Kγ inhibition. To test this possibility, we first validated the upregulation of these five genes upon eganelisib, but not LY294002 treatment, in CAL-1 and DF-DCN-1 cells (**Extended Data Fig. 6c**). Then, we depleted each gene individually in CAL-1 cells and treated them with eganelisib. Knockdown of 4 out of the 5 genes reduced sensitivity to PI3Kγ inhibition (**Extended Data Fig. 6d,e**). To validate the involvement of NFκB in activation of these tumor suppressor genes, we treated CAL-1 cells with selective NFκB inhibitors, IKK-16 and SC-514, and found that eganelisib no longer increased expression of those 5 genes (**Fig. 4f and Extended Data Fig. 6f**). Metabolic assays showed that NFκB inhibition blocked suppression of OXPHOS upon PI3Kγ inhibitor treatment (**Fig. 4g**). Finally, CAL-1 cells treated with NFκB inhibitors were rendered resistant to eganelisib (**Fig. 4h**). Mild-moderate growth suppressive effects of NFκB inhibitors on extended leukemia cell growth is not surprising considering the complexity of NFκB activities and is consistent with previous observations of susceptibility in BPDCN^39^ (**Extended Data Fig. 6g**). Together, these data support a conclusion that the PI3Kγ dependency is at least partially mediated by an NFκB-related transcriptional network.

### PI3Kγ inhibition suppresses a noncanonical PI3K pathway that is not dependent on Akt kinase (AKT)

We next focused on elucidating the substrate(s) of PI3Kγ in cells with activated PIK3R5/PIK3CG. As the most well-defined substrate of PI3K, AKT mediates nearly all the described functions of PI3K complexes, including PI3Kγ^14,40^. To our surprise, as measured by phosphoproteomics, AKT phosphorylation was not significantly changed in CAL-1 cells after PI3Kγ inhibition (**Fig. 5a**). GSEA also showed that the canonical PI3K-AKT-mTOR signaling target genes were not altered either after eganelisib treatment or PIK3R5 depletion (**Extended Data Fig. 7a**). These results indicated that noncanonical PI3K mediators may contribute to the dependency of the PIK3R5-related leukemia subset on PI3Kγ. Upon further analysis of proteomics and phosphoproteomics data, we found the phosphorylation of several proteins were affected by both PI3Kγ inhibition and PIK3R5 depletion (**Fig. 5a**). We focused on PAK1 for further study because PAK family members play important roles in oncogenic signal transduction including in leukemia^41,42^, PAK1 has not previously been connected with PI3Kγ, and there were validated PAK1 inhibitors and phospho-specific antibodies available. Western blotting confirmed that phosphorylation of PAK1 at serine 144 (S144), but not phosphorylation of other known proteins in canonical PI3K signaling, was significantly reduced upon PIK3R5/PIK3CG depletion (**Fig. 5b**). Moreover, treatment with eganelisib, but not LY294002, reduced PAK1 S144 phosphorylation in two independent cell lines (**Fig. 5c**). Of note, depletion of PIK3R5 or PIK3CG led to a complementary increase of AKT phosphorylation (**Fig. 5b**), whereas eganelisib induced suppression of AKT phosphorylation (**Fig. 5c**). This is consistent with the known activity of eganelisib on the suppression of AKT phosphorylation possibly via its inhibition of class IA PI3Ks^43,44^, which is why we focused on shared events seen after eganelisib treatment and PIK3R5 knockdown to provide specificity.

**Figure 5.**
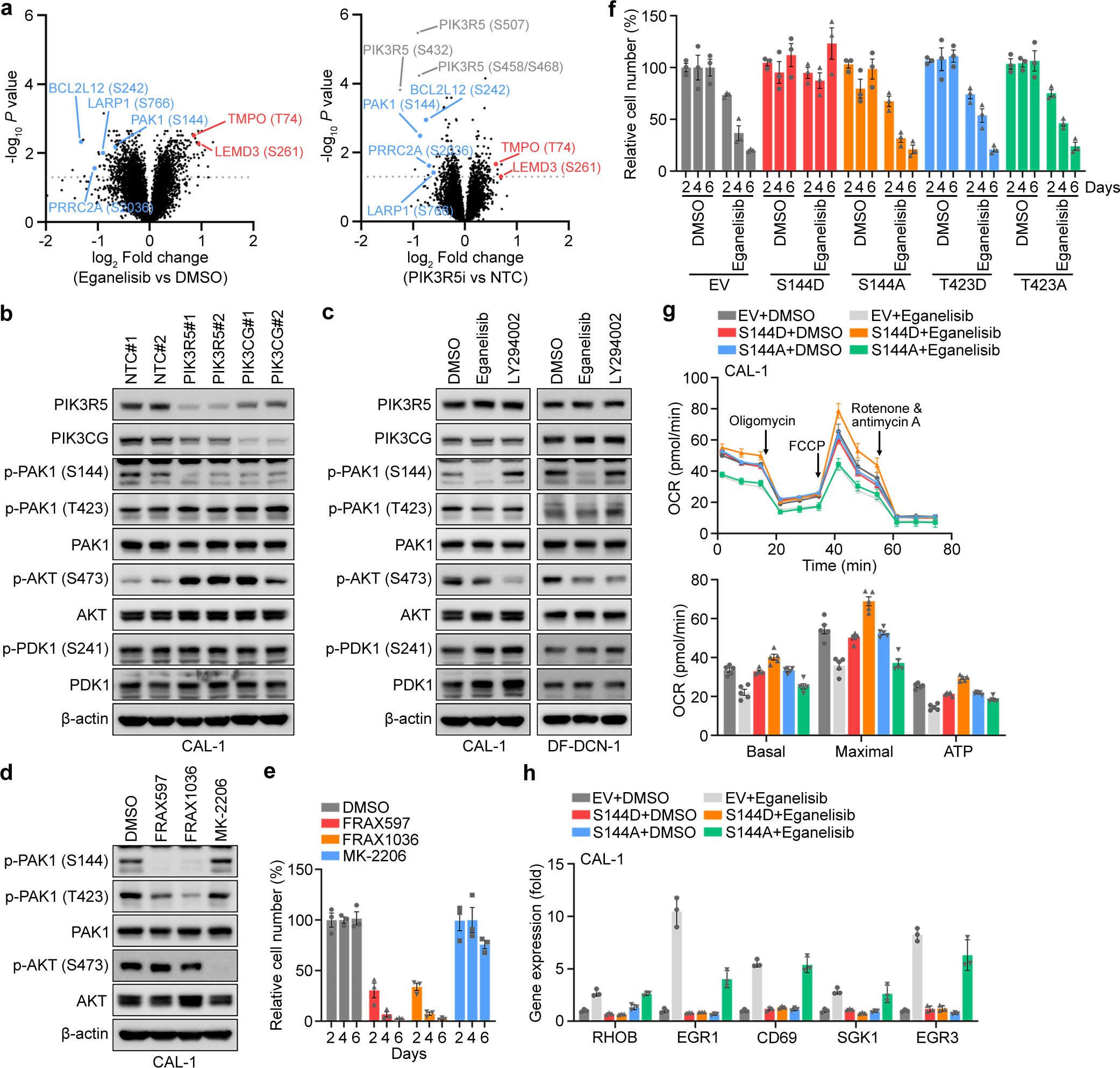
The intrinsic dependency of leukemia cells on the PI3Kγ complex is reliant on PAK1 but not AKT. **a,** Volcano plot showing consistently upregulated (red) or downregulated (blue) phosphorylation sites in CAL-1 cells treated with eganelisib or depleted of PIK3R5 (PIK3R5i). Adjusted *P* values are shown. **b,** Western blotting for the indicated proteins in control, PIK3R5 depleted, or PIK3CG depleted CAL-1 cells. **c,** Western blotting for the indicated proteins in CAL-1 and DF-DCN-1 cells after treatment with eganelisib or LY294002 (each 1 μM for 48 hours). **d,** Western blotting for the indicated proteins in CAL-1 cells upon treatment with FRAX597, FRAX1036, or MK-2206 (each 1 μM for 48 hours). **e,** Relative numbers of CAL-1 cells upon treatment with FRAX597, FRAX1036, or MK-2206 (each 1 μM) for 2, 4, and 6 days. Data are means ± S.E.M. from 3 biological replicates. **f,** Relative cell numbers of CAL-1 cells expressing empty vector (EV) or PAK1 mutants after treatment with vehicle control (DMSO) or eganelisib (1 μM) for 2, 4, and 6 days. Data are means ± S.E.M. from 3 biological replicates. **g,** Seahorse assays detecting mitochondrial respiration in CAL-1 cells expressing empty vector (EV) or PAK1 mutants after treatment with vehicle control (DMSO) or eganelisib (1 μM) for 2 days. Data are means ± S.D. from 5 technical replicates. **h,** RT-qPCR detection of gene expressions in CAL-1 cells expressing empty vector (EV) or PAK1 mutants after treatment with vehicle control (DMSO) or eganelisib (1 μM) for 2 days. Data are means ± S.D. from 3 technical replicates.

To further validate the functional involvement of PAK1 S144 phosphorylation in PI3Kγ dependency, we treated CAL-1 cells with selective inhibitors of PAK1 or AKT. PAK1 inhibitor treatment mimicked the strong suppression of CAL-1 growth that was observed with PI3Kγ inhibition, while the AKT inhibitor MK-2206 only minimally impaired cell viability despite completely abrogating AKT phosphorylation (**Fig. 5d,e**). Then, we asked if S144 is the critical phosphorylation site that mediates PAK1 function downstream of PI3Kγ. We introduced exogenous constitutively active (S144D) or inactive (S144A) PAK1 phospho-site mutants into CAL-1 cells and treated them with eganelisib. The introduction of active PAK1^S144D^, but not inactive PAK1^S144A^, conferred resistance to PI3Kγ inhibition (**Fig. 5f and Extended Data Fig. 7c**). As controls, introduction of either active of inactive PAK1 mutants at threonine 423 (T423)^45^ showed no effect on sensitivity to PI3Kγ inhibition (**Fig. 5f and Extended Data Fig. 7c**). Last, we tested the effect of PAK1 S144 phosphorylation on PI3Kγ downstream events. Seahorse assays indicated that introduction of PAK1^S144D^ abolished the suppression of OXPHOS by PI3Kγ inhibition (**Fig. 5g**). Eganelisib-induced elevation of the NFκB-related tumor suppressor genes was also diminished in CAL-1 cells expressing PAK1^S144D^ (**Fig. 5h**). Taken together, these results indicated that PAK1 is a critical substrate of PI3Kγ in the leukemia-intrinsic dependency pathway and phosphorylation of PAK1 S144 plays the essential role in this signaling.

### PI3Kγ inhibition synergizes with cytarabine in leukemia treatment

To explore the translational potential of targeting PI3Kγ in leukemia, we first tested the combination of eganelisib with cytarabine using two BPDCN cell lines with elevated PIK3R5. The rationale for combining with cytarabine includes: 1. Cytarabine is among the most commonly used chemotherapeutic drugs across acute leukemia histologies (in AML, ALL, and BPDCN)^46^; 2. Cytarabine-persistent leukemia cells exhibit elevated GPCR signaling, which is an upstream signal known to activate PI3Kγ^47–49^; 3. Cytarabine-persistent leukemia cells have increased dependence on OXPHOS^50^. As predicted, we observed significant synergistic effects between eganelisib and cytarabine in both cell lines (**Extended Data Fig. 8a**). Next, we utilized the intradermal xenografting model of DF-DCN-86 (**Extended Data Fig. 2c,d**) to assess the dynamic response of leukemia cells to eganelisib and cytarabine in vivo, alone or in combination. We found that single agent eganelisib was more effective than cytarabine alone, and the combination of these two drugs generated a synergistic suppressive effect on tumor growth (**Fig. 6a**). Moreover, western blotting of tumor lysates harvested immediately after treatment showed reduction of PAK1 S144, but not T423, phosphorylation, in animals receiving eganelisib or eganelisib plus cytarabine (**Fig. 6b and Extended Data Fig. 8b**). Last, we measured the effect of PI3Kγ inhibition on survival in six disseminated PDX models, three each PIK3R5-high or PIK3R5-low. As predicted, single agent eganelisib significantly prolonged survival in AML, ALL, and BPDCN with high PIK3R5, while having no benefit in cases with low PIK3R5 (**Fig. 6c**). However, to our surprise, the combination of eganelisib and cytarabine substantially increased survival in all cases compared to single agent cytarabine, regardless of baseline PIK3R5 (**Fig. 6c**).

**Figure 6.**
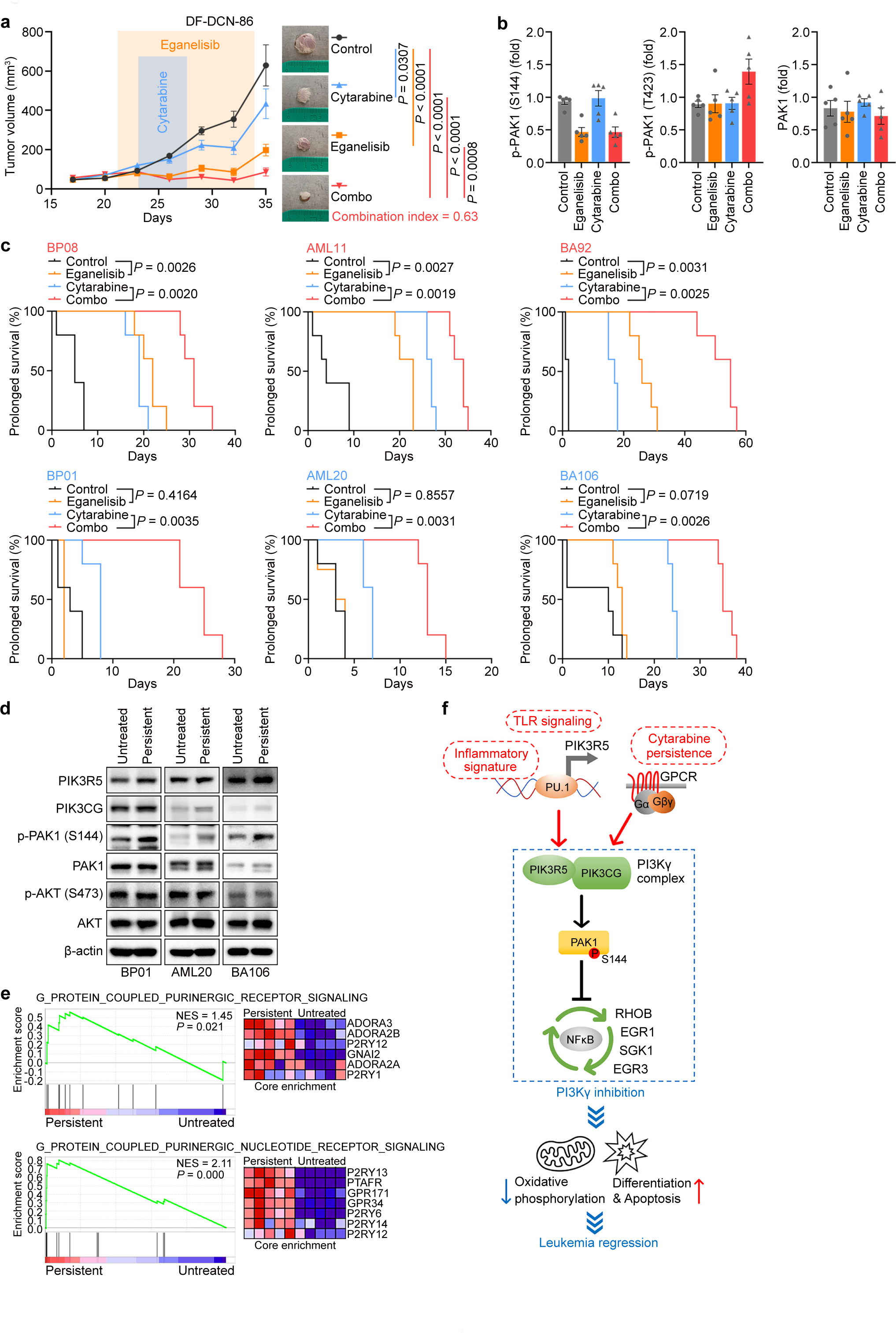
Eganelisib synergizes with cytarabine in leukemia treatment. **a,** Tumor volume and representative images of BPDCN intradermal xenografts. Data are means ± S.E.M. from 5 biological replicates. Two-way ANOVA test. **b,** Quantitative analysis of p-PAK1 and PAK1 expression levels in BPDCN intradermal xenografts. Data are means ± S.E.M. from 5 biological replicates. **c.** Survival of mice xenografted with leukemia PDXs and treated with eganelisib, cytarabine, or both (combo). Log-rank test. PIK3R5-high: BP08, AML11, and BA92; PIK3R5-low: BP01, AML20, and BA106. **d,** Western blotting for the indicated proteins in untreated or cytarabine-persistent PDX cells (BP01: 320 μM; AML20: 5 μM; BA106: 10 μM; each for 72 hours). **e,** GSEA indicating G protein-coupled purinergic receptor signaling pathways are significantly elevated in cytarabine-persistent AML20 cells. **f,** Proposed model illustrating leukemia dependency on noncanonical PI3Kγ signaling. The PI3Kγ complex can be activated via two mechanisms: 1. Intrinsically (baseline inflammatory signature) or extrinsically (via TLRs) activated innate inflammatory signaling promotes high PIK3R5 expression and subsequent stabilization of the PI3Kγ complex; 2. Leukemias without baseline elevated PIK3R5 acquire increased PI3Kγ activity through G protein-coupled purinergic receptor signaling after cytarabine treatment. In either circumstance, PI3Kγ activation leads to increased PAK1 phosphorylation, which drives leukemia via suppression of an NFκB-mediated transcriptional network of tumor suppressor genes. Inhibition of PI3Kγ reactivates the NFκB-related tumor suppressor genes, leading to compromised OXPHOS and leukemia cell death.

Based on the in vivo efficacy, we asked why the combination of eganelisib and cytarabine was synergistic even in leukemias without elevated PIK3R5. To begin, we performed ex vivo treatment of the three PDXs with low PIK3R5 expression at cytarabine concentrations resulting in ∼20% persistent leukemia cells, to enable analysis of residual disease (**Extended Data Fig. 8c**). Strikingly, the cytarabine-persistent cells exhibited substantially elevated phosphorylation of PAK1 S144, but not AKT S473, providing a potential explanation for their induced eganelisib sensitivity (**Fig. 6d**). There was a moderate and inconsistent increase of PIK3R5 or PIK3CG in some cytarabine-persistent leukemias (**Fig. 6d**), but this did not seem sufficient to explain the striking increase of PAK1 phosphorylation and the observed in vivo synergy between cytarabine and eganelisib. Thus, we asked if there were other potential mechanisms contributing to PAK1 activation. We established an in vivo cytarabine-persistent residual disease model using the AML20 PDX (**Extended Data Fig. 8d,e**). GSEA of RNA-seq comparing cytarabine-persistent to untreated cells indicated that G protein-coupled purinergic receptor signaling, but not other GPCR or PI3K-AKT-mTOR signaling, was activated in the cytarabine-persistent population (**Fig. 6e and Extended Data Fig. 8f**). Intriguingly, the G protein-coupled purinergic receptor family was recently found to activate PI3Kγ, but not other PI3K complexes, in the setting of leukemia resistance to targeted therapy^40^. Thus, our data suggest that activated G protein-coupled purinergic receptor-PI3Kγ-PAK1 signaling in cytarabine-persistent residual leukemia may also be targeted by PI3Kγ inhibition, expanding the potential clinical application of PI3Kγ inhibitors to leukemias without baseline PIK3R5 activation (**Fig. 6f**).

## Discussion

PI3Kγ has been pursued as a target in solid tumors because its suppression stimulates antitumor immune responses via reprograming tumor-associated macrophages^12,14^. The PI3Kγ inhibitor eganelisib is currently in Phase II clinical trials for solid cancers (e.g., NCT03961698 and NCT03980041). PI3Kγ inhibition appears to be well tolerated, supporting its potential expansion to other malignancies. However, targeting PI3Kγ has not had a direct effect on solid tumor cells in studies to date^12^, and thus it has been unclear whether any cancers have a cell-intrinsic dependency on this complex. We found that a leukemia subset characterized by innate inflammatory signaling-activated *PIK3R5* is highly sensitive to PI3Kγ inhibition. Surprisingly, expression of the PI3Kγ enzymatic subunit gene *PIK3CG* was not elevated, but its corresponding protein was increased. We determined that the PIK3R5 and PIK3CG proteins protect each other from UPS-mediated degradation, thus forming a self-stabilizing complex capable of transmitting even moderate transcriptional elevation of *PIK3R5* into substantial posttranslational activation of the PI3Kγ complex.

We found that PU.1 is the transcription factor mediating *PIK3R5* activation by inflammatory signaling. PU.1 is important for normal hematopoietic development^26,27^ and modulating PU.1 can promote myeloid differentiation in certain contexts^51–53^. In some leukemia cells, inhibition of PU.1 is an effective treatment^30,31^, but a systemic therapy targeted at PU.1 might induce severe toxicity to normal hematopoiesis. Here, we provide evidence of a functional axis of inflammatory signaling-PU.1-PI3Kγ that activates a survival pathway in some leukemias. This highlights a potential advantage to targeting PI3Kγ, rather than PU.1 or another upstream node, as this may preferentially affect PIK3R5-high leukemias but spare normal tissues without *PIK3R5* activation.

Unexpectedly, the intrinsic dependency of leukemias on PI3Kγ is not related to the conventional PI3K substrate AKT^14,40^ but rather relies on the phosphorylation of PAK1 kinase. PAK1 is involved in MAPK/ERK^54–56^ and canonical PI3K/AKT^57,58^ signaling, but has not been linked to PI3Kγ directly. Of potential relevance, PAK1 was found to mediate resistance to the PI3K inhibitors BEZ235 and BGT226 in lymphoma, which both target multiple PI3K complexes including PI3Kγ, but the downstream mechanism was not defined^59^. Through mass spectrometry phosphoproteomics, we identified PAK1 S144 to be one of the consistently downregulated phosphorylation sites upon both PIK3R5 knockdown and eganelisib treatment. Several ATP competitive and non-competitive allosteric inhibitors of PAK are available, and some have been tested in early phase clinical trials^60^. Future work will be required to determine the relative contribution of PAK1 and other potential PI3Kγ substrates in mediating leukemogenesis and therapeutic sensitivity in the PIK3R5-related leukemia subset. Additionally, mechanisms of resistance to PI3Kγ inhibition and whether resistant cells would remain dependent on downstream PAK1 are unknown. PAK1 inhibitors may help address these questions and provide another therapeutic node to target the leukemia intrinsic PI3Kγ pathway.

Cytarabine is a component of most conventional chemotherapy regimens used in AML, ALL, and BPDCN. Previous studies revealed that cytarabine-resistant leukemia cells have elevated OXPHOS^50^ and GPCR signaling^47^, which, given the consequences of PI3Kγ inhibition we observed here, led us to combine eganelisib and cytarabine. We found that this combination exerted synergy in models with elevated PIK3R5, as predicted. The combination was also effective in leukemias with low baseline PIK3R5, possibly related to G protein-coupled purinergic receptor signaling and PAK1 phosphorylation in cytarabine-persistent leukemia cells. Given that many patients with leukemia achieve remission after initial therapy yet nearly all will relapse without additional consolidation treatment, there has been increasing focus on targeting residual chemotherapy-persistent leukemia initiating cells or leukemia stem cells (LSCs). Important next steps from this work include optimization of timing to apply PI3Kγ inhibition for maximum clinical benefit, such as concurrently or sequentially with conventional chemotherapy, and whether similar PI3Kγ dependency is induced by therapies other than cytarabine.

Notably, a recent study identified a novel type of monocytic LSCs associated with clinical resistance to the BCL-2 inhibitor venetoclax^61^, and leukemias with monocytic pathologic features have poor outcomes with venetoclax based treatment regimens^62^. This is particularly interesting because our data suggested a monocytic lineage bias of the PIK3R5-related, eganelisib sensitive AML subset. Therefore, additional future work should investigate whether there is a synergistic effect between PI3Kγ inhibition and BCL-2 inhibition. The reported monocytic LSCs were also highly reliant on purine metabolism and targetable by cladribine, a purine analog that inhibits DNA/RNA synthesis^61,63^. Given that we found G protein-coupled purinergic receptors were activated in the cytarabine-persistent population, follow up studies should explore mechanistic interactions between purine metabolism and G protein-coupled purinergic nucleotide receptor signaling, including how they may increase dependence on PI3Kγ activity.

## Materials and methods

### Cell culture

293T, U-937, and THP-1 cells were purchased from ATCC (Manassas, VA). P31-FUJ and MV-4-11 cells were from Dr. James Griffin (Dana-Farber Cancer Institute [DFCI], Boston, MA). CAL-1 cells were provided by Dr. Takahiro Maeda (Nagasaki University, Nagasaki, Japan). 293T cells were cultured in DMEM supplemented with 10% fetal bovine serum (FBS) (F2442; Sigma-Aldrich, St. Louis, MO). U937, P31-FUJ, and CAL-1 cells were cultured in RPMI 1640 supplemented with 10% FBS (R10). THP-1 cells were cultured in R10 supplemented with 0.05 mM 2-mercaptoethanol (21985023; Thermo Fisher Scientific, Waltham, MA). MV-4-11 cells were cultured in IMDM supplemented with 10% FBS. Cell line identity was verified by short tandem repeat (STR) profiling in the Molecular Diagnostics Laboratory at DFCI (**Extended Data Table 1**), and cells were verified to be Mycoplasma-free by regular testing.

The BPDCN cell line DF-DCN-86 was established from a skin biopsy in a BPDCN patient. Briefly, the skin biopsy sample was cut into small pieces with a scalpel then digested with type II collagenase (17101015; Thermo Fisher Scientific) at 37 °C for 30 min. After filtering with a 70 µm cell strainer (352350; Corning, Glendale, AZ), the cells were washed once with 1X RBC lysis buffer (420301; BioLegend, San Diego, CA) then cultured with advanced DMEM/F-12 (12634028; Thermo Fisher Scientific) based leukemia adaptive medium (LAM) for 2 weeks: 1X Penicillin-Streptomycin (15140122; Thermo Fisher Scientific), 100 µg/ml Primocin (ant-pm-1; InvivoGen, San Diego, CA), 0.5 µg/ml Caspofungin (SML0425; Sigma-Aldrich), 1X HEPES (15630080; Thermo Fisher Scientific), 1X Glutamax (35050061; Thermo Fisher Scientific), 10 µM Y27632 (130-106-538; Miltenyi Biotec, Bergisch Gladbach, North Rhine-Westphalia, Germany), 100 ng/ml R-spondin-1 (120-38; PeproTech, Cranbury, NJ), 100 ng/ml Noggin (120-10C; PeproTech), 1X B-27 (17504044; Thermo Fisher Scientific), 500 nM A 83-01 (SML0788; Sigma-Aldrich), 1 µM Prostaglandin E_2_ (2296; Tocris Bioscience, Bristol, UK), 1.25 mM *N*-Acetyl-L-cysteine (A8199; Sigma-Aldrich), 10 mM Nicotinamide (N0636; Sigma-Aldrich), 3 µM CHIR99021 (SML1046; Sigma-Aldrich), 10 µM SB 202190 (S7076; Sigma-Aldrich), 50 ng/ml EGF (AF-100-15; PeproTech), 10 ng/ml FGF-10 (100-26; PeproTech), 10 ng/ml FGF-basic (100-18B; PeproTech), 50 ng/ml Flt3-Ligand (300-19; PeproTech), 10 ng/ml SCF (300-07; PeproTech), 10 ng/ml IL-3 (200-03; PeproTech), and 1 µM StemRegenin 1 (S2858; Selleckchem, Houston, TX). Then, the LAM was replaced with R10 by 10% increasing each week for 9 weeks until the culture contained 10% LAM and 90% R10, after which the percentage of LAM was further reduced to 5%, 2%, 1% and 0% each week. The DF-DCN-1 and DF-DCN-2 cell lines were established by a similar method from BPDCN primary bone marrow samples that were previously grown in vitro in coculture with MS-5 stromal cells. Those cells were then cultured with the mix of LAM and R10 (supplemented with Flt3-Ligand, SCF, IL3 and SR-1 as the same concentrations as in LAM) at a ratio of 1:1, and the percentage of LAM was further reduced to 40%, 30%, 20%, 10%, 5%, 2%, 1% and 0% each week. All the 3 cell lines were cryopreserved in FBS + 10% DMSO, and 10% LAM was supplemented in the culture medium during the first 2 passages after each freeze-thaw cycle or after lentivirus infection.

### Reagents

Eganelisib (S8330), duvelisib (S7028), LY294002 (S1105), resiquimod (S8133), IKK-16 (S2882), SC-514 (S4907), FRAX597 (S7271), FRAX1036 (S7989), MK-2206 (S1078), cytarabine (S1648), and MG132 (S2619) were purchased from Selleckchem for in vitro and ex vivo experiments. Eganelisib (HY-100716) and cytarabine (HY-13605) were purchased from MedChemExpress (Monmouth Junction, NJ) for in vivo experiments. Emetine dihydrochloride (sc-202600) was purchased from Santa Cruz Biotechnology (Dallas, TX).

### Plasmids

Guide RNA sequences for CRISPRi were cloned into pXPR_050 vector (#96925; Addgene, Watertown, MA). shRNA sequences were cloned into pSIH1-H1-puro vector (#26597; Addgene). cDNA sequences of PAK1 mutants with a C-terminus Flag tag were synthesized at BGI Genomics (Beijing, China) then cloned into pLVX-IRES-Neo vector (632184; Takara Bio, Kusatsu, Shiga, Japan). All sequences are in **Extended Data Table 2**.

### Analyses of TCGA AML

All patients in the TCGA AML dataset with both RNA-seq and prognosis data available were included for overall survival analyses (n=161). The overall survival was calculated from the date of first treatment until death or last follow-up and comparisons were made by the Log-rank test. Sixteen patients with *PML-RARA* rearrangement were excluded from the dataset before regression analyses as those patients are clearly distinguished (belonging to French-American-British M3 pathologic subtype) from other AML patients and have a long-term survival rate up to 90%. Categorical variables were summarized as counts and percentages, and comparisons were made by Pearson’s chi-square test. Continuous variables were summarized as median and range, and comparisons were made by the Wilcoxon rank sum test. Cox proportional hazards regression model was fit in a stepwise backward selection to assess the effect of the IIRS score adjusted for age and disease characteristics on OS. Log-rank test was performed by GraphPad Prism 9 (GraphPad Software, San Diego, CA), and other analyses were performed by Stata 18.0. (StataCorp, College Station, TX).

### Virus production and transduction

Lentivirus was produced by transfection of 293T cells with the second-generation packaging system psPAX2 (#12260, Addgene) and pMD2.G (#12259, Addgene). 293T cells were seeded at a confluence of 95% and culture medium was replaced with fresh DMEM without serum or antibiotics before transfection. Transfections were performed using Lipofectamine 2000 Transfection Reagent (11668500, Thermo Fisher Scientific) according to the manufacture’s protocol. Six hours after transfection, the medium was replaced with regular DMEM culture medium that contains 10% FBS and 1X Penicillin-Streptomycin. Viral supernatants were collected 48 and 72 hours after transfection and filtered through a 0.45 μm filter. Lentivirus transduction was performed by spin infection. Briefly, 2-5 × 10^5^ cells were resuspended in 500 μl culture medium and seeded into a well of 12-well plate, then 1 ml lentivirus was added and Polybrene Infection/Transfection Reagent (TR-1003-G; Sigma-Aldrich) was supplemented at a final concentration of 8 μg/ml. After spinning at 930 g, 30 °C for 2 hours with zero brake, another 1 ml regular culture medium was slowly added to each well without disturbing the cells. One day after the spin infection, virus-containing medium was replaced with fresh culture medium and cells were allowed to recover for another day, then cells were selected with 2 μg/ml puromycin (631306; Takara Bio) or 500 μg/ml geneticin (10131027; Thermo Fisher Scientific) for 3-7 days before use in experiments.

### CRISPR interference screening and analysis

The CRISPR interference screen was performed in the BPDCN cell line CAL-1, with two biological replicates. KRAB-dCas9 (#96918; Addgene) was transduced by lentivirus and successfully transduced cells were selected by 6 μg/ml blasticidin (R21001; Thermo Fisher Scientific) for 1-2 weeks. The expression of KRAB-dCas9 was validated by western blotting and a similar growth rate was observed between KRAB-dCas9-expressing and parental CAL-1 cells. The virus for the genome-wide Dolcetto CRISPR interference library^15^ was obtained from the Genetic Perturbation Platform of Broad Institute (Cambridge, MA), which includes Set A and Set B that contain 57,050 and 57,011 sgRNAs to target 18,901 and 18,899 genes, respectively. KRAB-dCas9-expressing CAL-1 cells were transduced with the Dolcetto library at a multiplicity of infection of 0.3 by spin infection. Two days after transduction, cells were selected with 2 μg/ml puromycin for another 6 days, then samples were collected as day 0 of the dependency screen. The remaining cells were passaged every 2-3 days for collection of day 14 and day 21 samples. The sgRNA coverage was >1,000x throughout the screen and sample collections. Genomic DNA was extracted using a NucleoSpin Blood XL kit (740950.50; Takara Bio) and further cleaned up by a OneStep PCR Inhibitor Removal Kit (D6030; Zymo Research, Irvine, CA) before PCR amplification. Sequencing and data analysis were performed by the Genetic Perturbation Platform of Broad Institute. Screening results are in **Extended Data Table 3**.

### Short-term cell proliferation assay

Cells were washed once with serum-free medium then cultured with the corresponding base medium supplemented with 0.5-5% FBS. The starting cell density (day 0) was 1 × 10^5^/ml and the cell numbers were counted by a Countess II Automatic Cell Counter (AMQAX1000; Thermo Fisher Scientific) every 2 days till day 6. Relative cell number (%) was calculated by the ratio of experimental arm (gene silencing or inhibitor treated) to control arm (nontargeting control or DMSO treated) at each time point.

### Ex vivo and in vitro drug sensitivity assays

For ex vivo drug sensitivity assays using PDXs, PDX cells were collected from the bone marrow and spleen of NOD.Cg-*Prkdc^scid^ Il2rg^tm1Wjl^*/SzJ (NSG; 005557; The Jackson Laboratory, Bar Harbor, ME) mice, then purified with a Mouse Cell Depletion Kit (130-104-694; Miltenyi Biotec) and/or Dead Cell Removal Kit (130-090-101; Miltenyi Biotec) to ensure the human cell percentage and live cell percentage were both greater than 90%. PDX cells were seeded into white 96-well plates (3917; Corning) with 4 × 10^4^ cells per well in 100 μl LAM, then treated for 72 hours. Cell viability was determined by a CellTiter-Glo 2.0 Cell Viability Assay (G9242; Promega, Madison, WI) according to the manufacture’s protocol. In vitro drug sensitivity assays using leukemia cell lines were performed similarly except for adjusting the cell number per well to 0.5-4 × 10^4^ according to the growth rates of different cell lines. Synergy score was calculated using the SynergyFinder online tool (https://tangsoftwarelab.shinyapps.io/synergyfinder/).

### Caspase-3/7 activity measurement

Caspase-3/7 activity in leukemia cells were detected by Caspase-Glo 3/7 Assay System (G8091; Promega) according to the manufacturer’s protocol. Briefly, PDX cells were seeded into white 96-well plates (3917; Corning) with 4 × 10^4^ cells per well in 100 μl LAM, then treated for 72 hours. Then, 100 μl premixed Caspase-Glo 3/7 reagent was added into each detection well. After mixing the contents on a plate shaker for 2 minutes, the plate was incubated at room temperature for 1 hour before being measured by a SpectraMax M3 Multi-Mode Microplate Reader.

### Intradermal xenografting model

The BPDCN cell line DF-DCN-86 established from a patient skin tumor was used for an intradermal xenografting model to mimic the clinical characteristics of BPDCN patients. Briefly, 1.25-10×10^6^ DF-DCN-86 cells were suspended in 50 μl cold PBS, then mixed with 50 μl Cultrex UltiMatrix Reduced Growth Factor Basement Membrane Extract (BME001-05; R&D Systems, Minneapolis, MN) on ice. Hair of NSG mice around the xenografting area was shaved with an electric clipper, and the 100 μl cell mixture was injected intradermally using a 28-gauge insulin syringe. Tumor size was measured by a caliper and tumor volume was calculated using the following formula: 0.52 × length × width^2^. After the tumors had grown for the designated time, mice were euthanized and the tumors were harvested for protein extraction and western blotting.

### Advanced stage systemic PDX model

To evaluate the effects of eganelisib alone or in combination with cytarabine on overall survival, 1 × 10^6^ leukemia PDX cells were suspended in PBS containing 0.2% FBS then injected into a NSG mouse via tail vein. Treatment was started 3 weeks post-xenografting when the disease was in an advanced stage (with a peripheral blood leukemia burden approximately 1-5%). Eganelisib was given at 15 mg/kg daily for 14 days via oral gavage (dissolved in 10% DMSO, 40% PEG300, 5% Tween-80, and 45% saline sequentially), and cytarabine was given at 30 mg/kg daily for 5 days (day 3-7 of eganelisib treatment period) via intraperitoneal injection (dissolved in saline). The date when the first mouse in control group reached the endpoint was recorded as the start day for the assessment of prolonged survival. Mice were humanely sacrificed once moribund.

### Residual disease leukemia model

To obtain a persistent leukemia population after cytarabine treatment, 1 × 10^6^ acute leukemia PDX cells were suspended in PBS containing 0.2% FBS then injected into a NSG mouse via tail vein, and treatment was started 3 weeks post-xenografting. Cytarabine was given at 30 mg/kg daily for 5 days via intraperitoneal injection. One day after the last dose of treatment, mice were sacrificed and cells were collected from bone marrow. Human leukemia cells were labelled with a FITC anti-human CD45 antibody (368508; BioLegend) then sorted using a BD FACS Aria II Cell Sorter.

### Genetic characterization of leukemia models

BPDCN cell lines (**Extended Data Table 1**) and PDXs^19^ were characterized using OncoPanel, a custom capture-based next-generation sequencing based assay for detection of single-nucleotide variants, insertions/deletions, copy number alterations, and structural variants across 282 cancer genes^64^ and/or a PCR amplification-based hematologic malignancy gene panel^65^.

### Reverse transcription quantitative real-time PCR (RT-qPCR)

Total RNA was extracted using NucleoSpin RNA Plus XS (740990.250; Takara Bio) according to the manufacturer’s protocol. First strand cDNA synthesis was performed with LunaScript RT SuperMix Kit (E3010L; New England Biolabs, Ipswich, MA), and 100 ng-1 μg total RNA was used for one 20 μl RT reaction. qPCR analysis was conducted on a QuantStudio 6 Flex Real-Time PCR System (4485697; Thermo Fisher Scientific) with Luna Universal qPCR Master Mix (M3003X; New England Biolabs). The qPCR primer sequences were as follows (5’-3’): GAPDH (Forward-TCGGAGTCAACGGATTTG; Reverse-CAACAATATCCACTTTACCAGAG), PIK3R5 (Forward-TGACATGCTACTCTACTACTG; Reverse-GGAGTGGATGAAGATCTCTG), PIK3CG (Forward-TCAGGACATCTGTGTTAAGG; Reverse-GCATCCCGGATATATTCAATG), SPI1 (Forward-AGCCATAGCGACCATTAC; Reverse-CTCCGTGAAGTTGTTCTC), ASH2L (Forward-CTTTTGGATCAGGACCTTAG; Reverse-CAGAAAACAAAGGGTCACTC), RHOB (Forward-CAAGGAGAGGGAAAAGAAAC; Reverse-ACTGCCCTTTATCAAAACTG), EGR1 (Forward-CAAAATAAGGAAGAGGGCTG; Reverse-CTACAACATTCCAACTCCTG), CD69 (Forward-CTACTCTTGCTGTCATTGATTC; Reverse-GTTCCTTTTTCAGTCCAACC), SGK1 (Forward-GAGAAGCATATTATGTCGGAC; Reverse-TCTGGAGATGGTAGAACAAC), and EGR3 (Forward-CAGAGAATGTAATGGACATCG; Reverse-CATGAGGCTAATGATGTTGTC).

### Western blotting

Cells were lysed with RIPA Lysis and Extraction Buffer (89901; Thermo Fisher Scientific) for 10 minutes on ice. After brief sonication, cell lysate was centrifuged for 10 minutes at 20,000 g, 4 °C, to remove insoluble debris. Protein concentrations were determined by Pierce BCA Protein Assay Kit (23227; Thermo Fisher Scientific) and samples were prepared by boiling at 99 °C for 10 minutes with Blue Loading Buffer (7722S; Cell Signaling Technology, Danvers, MA). Then, 5-20 μg of total protein was separated by NuPAGE 4-12% Bis-Tris Mini Protein Gels (NP0336BOX; Thermo Fisher Scientific) and transferred to Immobilon-P PVDF Membrane (IPVH00010; MilliporeSigma, Burlington, MA). Following transfer, the membrane was blocked with 5% non-fat milk (NC9022655; Fisher Scientific, Waltham, MA) or bovine serum albumin (9998S; Cell Signaling Technology) for 45 minutes at room temperature. Primary antibodies were incubated at 4 °C overnight and secondary antibodies were incubated at room temperature for 90 minutes. Proteins were detected by Clarity Western ECL Substrate (1705061; Bio-Rad, Hercules, CA) and images were taken by a ImageQuant LAS 4000 or a ImageQuant 800 (Cytiva, Marlborough, MA). The following primary antibodies were used for Western Blotting: anti-PIK3R5 (1:500; sc-390916; Santa Cruz Biotechnology), anti-PIK3CG (1:500; sc-166365; Santa Cruz Biotechnology), anti-ubiquitin (1:1000; 43124S; Cell Signaling Technology), anti-SPI1 (1:1000; 2258S; Cell Signaling Technology), anti-ASH2L (1:2000; ab176334; Abcam, Cambridge, UK), anti-p-PAK1-S144 (1:1000; 2606S; Cell Signaling Technology), anti-p-PAK1-T423 (1:1000; ab2477; Abcam), anti-PAK1 (1:1000; 2602S; Cell Signaling Technology), anti-p-AKT-S473 (1:2000; 4060S; Cell Signaling Technology), anti-AKT (1:1000; 4691S; Cell Signaling Technology), anti-p-PDK1-S241 (1:1000; 3061S; Cell Signaling Technology), anti-PDK1 (1:1000; 3062S; Cell Signaling Technology), and anti-β-actin (1:5000; A5441; Sigma-Aldrich). Horse anti-mouse IgG (7076S; Cell Signaling Technology) and goat anti-rabbit IgG (7074S; Cell Signaling Technology) secondary antibodies were used at a dilution of 1:2000.

### Immunoprecipitation and ubiquitination assay

Ubiquitination was measured using control, PIK3R5-knockdown, or PIK3CG-knockdown CAL-1 cells. Cells were treated with MG132 (50 μM) for 8 hours before harvesting, then 5 × 10^7^ cells were lysed in 4 ml Pierce IP Lysis Buffer (87788; Thermo Fisher Scientific) for 20 minutes on ice. After centrifuging for 10 minutes at 20,000 g, 4 °C, 90 μl of the supernatant was taken from each condition as input, and the remaining supernatant was incubated with 20 μg anti-PIK3R5 (sc-390916; Santa Cruz Biotechnology) or anti-PIK3CG (sc-166365; Santa Cruz Biotechnology) antibodies overnight with rotation. Then, 50 μl Pierce Protein A/G Magnetic Beads (88802; Thermo Fisher Scientific) was added to each supernatant/antibody mixture and incubated for 2 hours at room temperature with rotation. Last, the beads were washed twice with 1x Pierce TBS Tween 20 Buffer (28360; Thermo Fisher Scientific) before proteins were eluted into 150 μl 1x Blue Loading Buffer (7722S; Cell Signaling Technology) by boiling at 99 °C for 10 minutes.

### Flow cytometry

Leukemia cells were resuspended in flow buffer (PBS containing 2% FBS) and cell surface proteins were stained with the following antibodies (5 μl per test in 100 μl flow buffer) at 4 °C for 20 minutes: FITC anti-human CD45 (368508; BioLegend), FITC anti-human CD14 (325604; BioLegend), and APC/Cyanine7 anti-mouse/human CD11b (101226; BioLegend). After staining, cells were washed with 1 ml flow buffer once then resuspended in 200 μl flow buffer and measured using a CytoFLEX cytometer (Beckman Coulter, Jersey City, NJ).

### ATAC-seq

Genomic DNA was extracted using the Quick-DNA Microprep Kit (D3020; Zymo Research). ATAC-seq was performed as previously described^66^. Reads were mapped to the Genome assembly GRCh37, and peaks were visualized via the Integrative Genomics Viewer^67^.

### ChIP assays

ChIP assays were performed using the SimpleChIP Enzymatic Chromatin IP Kit (9003S; Cell Signaling Technology) according to the manufacturer’s protocol. The following primary antibodies were used for ChIP: anti-Pol II (1 μg per IP prep; 14958S; Cell Signaling Technology) and anti-PU.1 (1 μg per IP prep; 2266S; Cell Signaling Technology). qPCR analysis was conducted on a QuantStudio 6 Flex Real-Time PCR System (4485697; Thermo Fisher Scientific) with Luna Universal qPCR Master Mix (M3003X; New England Biolabs). The qPCR primer sequences for the detection of PU.1 binding region were as follows (5’-3’): pair #1 (Forward-TCGGAAGCGCGGCTTTG; Reverse-CGCAGTGAGAGAAGGAAGCA), and pair #2 (Forward-GAAGCGCGGCTTTGGC; Reverse-TCGCAGTGAGAGAAGGAAGC).

### RNA sequencing (RNA-seq)

RNA-seq was performed by the Molecular Biology Core Facilities at DFCI (http://mbcf.dfci.harvard.edu/). The raw FASTQ data were analyzed by the VIPER pipeline to generate gene expression data, as previously described^18^. Gene set enrichment analysis (GSEA) was performed with the GSEA software according to published guidelines^68,69^.

### Quantitative Tandem Mass Tag (TMT) proteomics and phosphoproteomics

To quantify proteins and phosphopeptides, the streamlined (SL)-TMT method described by Navarrete-Perea and collaborators^70^ was followed. CAL-1 cells were lysed in an 8M urea buffer (8M Urea, 75 mM NaCl, 50 mM HEPES pH 7.5) supplemented with EDTA-free protease and phosphatase inhibitors (Roche). The lysates were then clarified by centrifugation at 17,000 x g for 15 min at 4°C and quantified. To reduce and alkylate cysteines, 200 µg of protein was sequentially incubated with 5 mM TCEP for 30 mins, 14 mM iodoacetamide for 30 mins (in the dark), and 10 mM DTT for 15 mins. All reactions were carried out at RT. Next, proteins were chloroform-methanol precipitated and the pellet resuspended in 200 mM EPPS pH 8.5. Then, the protease LysC (Wako) was added at 1:100 (LysC:protein) ratio and incubated overnight at RT. The following day, samples were further digested for 5 hours at 37°C with trypsin at 1:75 (trypsin:protein) ratio. Both digestions were performed using an orbital shaker at 1,500 rpm. After digestion, samples were clarified by centrifugation at 17,000 x g for 10 min. The peptide concentration in the supernatant was quantified using a quantitative colorimetric peptide assay from Thermo Fisher Scientific (Cat. No. 23275). TMT labeling was carried out using the TMTpro-18plex kit from Thermo Fisher Scientific. Briefly, for each sample, 100 μg of peptides was brought to 1 μg/μl with 200 mM EPPS (pH 8.5), acetonitrile (ACN) was added to a final concentration of 30% followed by the addition of 200 μg of each TMT label. After 1 h of incubation at RT, the TMT labeling was quenched by adding 0.3% hydroxylamine (Sigma) for 15 min at RT. Labeled samples were combined, desalted using tC18 SepPak solid-phase extraction cartridges (100 mg, Waters) as described in the previous study^70^, and dried in a SpeedVac. Next, desalted peptides were subject to a phosphopeptide enrichment (“mini-phos”) using the High-Select Fe-NTA Phosphopeptide Enrichment Kit (Cat# A32992, Thermo Fisher Scientific) following manufactureŕs instructions. The unbound fraction and first two washes where pooled, dried to near dryness using vacuum centrifugation, and subsequently fractionated though basic pH reversed phase chromatography using a HPLC equipped with a 3.5 µm Zorbax 300 Extended-C18 column (Agilent). The fractions were first collected into a 96-well plate and then combined into 24 samples. Twelve of them were desalted following the C18 Stop and Go Extraction Tip (STAGE-Tip)^71^ and dried down in a SpeedVac. These samples correspond to the total proteome analysis. Eluted peptides from the “mini-phos” enrichment were desalted using STAGE-Tip and dried down in the SpeedVac before MS analysis. Each of the twelve fractions from the total proteome was analyzed once, while the sample containing the phosphopeptides from the “mini-phos” was analyzed twice. All samples were analyzed in an Orbitrap Fusion Lumos mass spectrometer equipped with a FAIMSpro module, operating in high-resolution MS2 (hrMS2) mode^72,73^. The mass spectrometer was coupled to a Proxeon NanoLC 1200 (Thermo Fisher Scientific) mounted with a 100 μm capillary column that was packed with 35 cm of Accucore 150 resin (2.6 μm, 150 Å; Thermo Fisher Scientific). Peptides were separated at 525 nL/min flow rate using two buffers as mobile phases (Buffer A: 5% acetonitrile, 0.1% formic acid and Buffer B: 95% acetonitrile, 0.1% formic acid). For both total and phospho TMT analysis, different MS parameters were employed, including gradient length, FAIMS compensation voltage (CV), MS1 orbitrap resolution, scan range, MS1 maximum injection time, automatic gain control (AGC), MS2 isolation window, higher-energy collision dissociation (HCD), Orbitrap MS2 resolution, and MS2 maximum injection time and AGC. For additional details about the specific parameters and settings used in the analyses, please refer to **Extended Data Table 4**.

A suite of in-house pipeline (GFY-Core Version 3.8, Harvard University) was used to obtain final protein quantifications from all RAW files collected from the Orbitrap Fusion Lumos. RAW data were converted to mzXML format using a modified version of ReAdW.exe. mzXML files were searched using the search engine Comet^74^ against a human target-decoy protein database (downloaded from UniProt in April 2021) that included the most common contaminants^75,76^. Precursor ion tolerance was set at 20 ppm and product ion tolerance at 0.02 Da. TMTpro tags on lysine residues and peptide N termini (+304.2071 Da) and carbamidomethylation of cysteine residues (+57.021 Da) were set as static modifications, while oxidation of methionine residues (+15.995 Da) was set as a variable modification. For phosphorylation analysis, phosphorylation (+79.966 Da) on serine, threonine, and tyrosine were set as variable modifications. Peptide-spectrum matches (PSMs) were adjusted to a 1% FDR using a linear discriminant analysis as described previously^77^. For total proteome analysis, proteins were further collapsed to a final protein-level FDR of 1%. Phosphorylation site localization was determined using the AScore algorithm^78^. AScore is a probability-based method designed for the high-throughput localization of protein phosphorylation sites. A threshold of 13 corresponds to a 95% confidence level in site localization. TMT quantitative values we obtained from MS2 scans. Only those PSMs with a summed signal-to-noise ratio (S/N) across all samples > 100 and an isolation specificity > 0.7 were used for quantification. These PSMs were summed for each protein to obtain their relative abundance. To account for equal protein loading in both TMT (total and phosphor), all TMT intensities were normalized so the sum of the TMT signal for all proteins quantified in each channel was equivalent (**Extended Data Table 4**). Quantifications are represented as relative abundances (**Extended Data Table 5**).

### Metabolic phenotyping (Seahorse) assay

OXPHOS and glycolysis of leukemia cells were assessed by extracellular flux using a Seahorse XFe96 Analyzer (Agilent Technologies, Santa Clara, CA) with a Cell Mito Stress Test Kit (103015; Agilent Technologies) and a Glycolysis Stress Test Kit (103020; Agilent Technologies), respectively. XF RPMI Base Medium supplemented with 1 mM pyruvate, 2 mM glutamine, and 10 mM glucose (for the Cell Mito Stress Test Kit) or 1 mM glutamine (for the Glycolysis Stress Test Kit) was used as the assay medium (103681-100; Agilent Technologies). Briefly, Seahorse XF96 Cell Culture Microplates (101085; Agilent Technologies) were treated with Cell-Tak Cell and Tissue Adhesive (354240; Corning) for 20 minutes (22.4 μg/ml; 25 μl per well), after which each well was washed with 200 μl sterile water twice. Then, 2 × 10^4^ (CAL-1) or 5 × 10^4^ (DF-DCN-86 or DF-DCN-1) leukemia cells were resuspended in 50 μl of assay medium and plated into each well. The microplates were centrifuged at 200 g for 1 minute with zero braking, then transferred to a 37°C non-CO_2_ incubator for 25-30 minutes. Subsequently, another 130 μl of assay medium was added into each well and the microplate was incubated in a non-CO_2_ incubator for 15-25 minutes before being loaded into the XFe96 Analyzer. For the mitochondrial stress test, oligomycin, carbonyl cyanide-p-trifluoromethoxyphenylhydrazone (FCCP) and rotenone/antimycin A (Rot/AA) were sequentially injected to final concentrations of 1.5 μM, 0.5 (CAL-1) or 1.0 (DF-DCN-86 or DF-DCN-1) μM, and 0.5 μM, respectively. For the glycolysis test, glucose, oligomycin and 2-deoxy-D-glucose (2-DG) were sequentially injected to final concentrations of 10 mM, 1 μM, and 50 mM, respectively. Five replicates were included in each group, and data analyses were performed using Wave software (Agilent Technologies).

### Statistical analysis

Two-tailed *t* test, Mann-Whitney test, two-way ANOVA test, or Log-rank test were used for statistical analyses. For all statistical analyses, differences for which *P* ≤ 0.05 were considered statistically significant, and at least three biologically independent experiments with similar results were reported. The data are presented as means ± S.E.M. or means ± S.D. as indicated in the figure legends. The sample size (n) for each analysis was determined on the basis of pretests and previous similar experiments, and are indicated in the figure legends. Unless otherwhere stated, all other data analyses were performed using GraphPad Prism 9 (GraphPad Software).

## Acknowledgements

Q.L. is a Fellow of The Leukemia & Lymphoma Society. X.W. is supported by a National Cancer Center fellowship grant. C.A.G.B. is a Special Fellow of The Leukemia & Lymphoma Society. R.X. is a Dr. Oliver Press Memorial Fellow of Lymphoma Research Foundation. A.A.L. is a Scholar of The Leukemia & Lymphoma Society. A.A.L. is supported by the NCI (R37 CA225191), Department of Defense (W81XWH-20-1-0683), the Mark Foundation for Cancer Research, the Ludwig Center at Harvard, and the Bertarelli Rare Cancers Fund. None of the funding bodies were directly involved in the design of the study, nor in the collection, analysis, or interpretation of the data, or writing of the manuscript.

## Author contributions

Q.L., E.G.R., X.W., K.Y., C.A.G.B., and R.X. conducted most of the experiments and analyzed data. E.G.R., X.W., K.G., P.v.G., J.G.D., and D.S.N. assisted in experimental design and data interpretation. M.A.P. and J.A.P. performed proteomic and phosphoproteomic experiments and data analysis. S.S. conducted clinical data analysis. H.W.L. performed ATAC-seq analysis. Q.L. and A.A.L. designed the study, interpreted the data, and wrote the paper. A.A.L. supervised the study. All authors read and participated in editing the paper.

## Data availability

The RNA-seq data are available at the Gene Expression Omnibus under accession numbers GSE243654 (PI3Kγ inhibition in CAL-1 cells) and GSE243669 (cytarabine persistent leukemias). The ATAC-seq data are available at the Gene Expression Omnibus under accession number GSE243666. The mass spectrometry data have been deposited to the ProteomeXchange Consortium via the PRIDE partner repository^79^ with the dataset identifier PXD045702.

## Extended Data Tables

**Extended Data Table 1**. Genomic profiling of BPDCN cell lines.

**Extended Data Table 2**. Fold change of all gene targets in CRISPRi screening.

**Extended Data Table 3**. Sequences of CRISPRi, shRNA, and cDNA.

**Extended Data Table 4**. Mass spectrometry parameters.

**Extended Data Table 5**. Protein and phosphoprotein quantification.

---

**Extended Data Figure 1.**
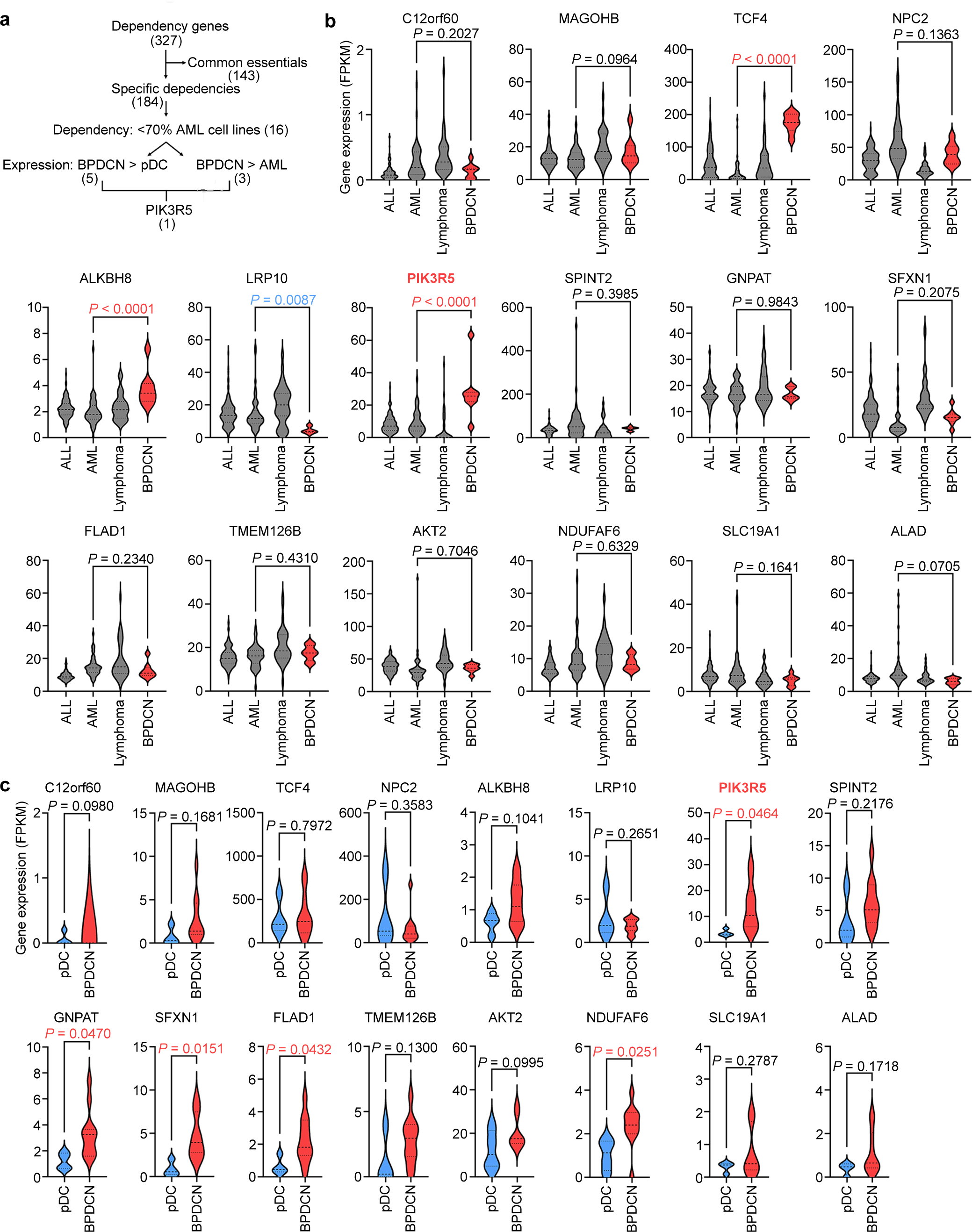
PIK3R5 expression is significantly elevated in BPDCN. **a,** Flowchart showing the bioinformatic screening strategy to identify potential specific targets in BPDCN. **b,** Expression of the 16 genes that are dependencies in <70% of AML cell lines from panel a in various blood cancer subtypes. Two-tailed *t* test. **c,** Expression of the 16 genes that are dependent in <70% of AML cell lines from panel a in normal pDC and BPDCN. Two-tailed *t* test.

**Extended Data Figure 2.**
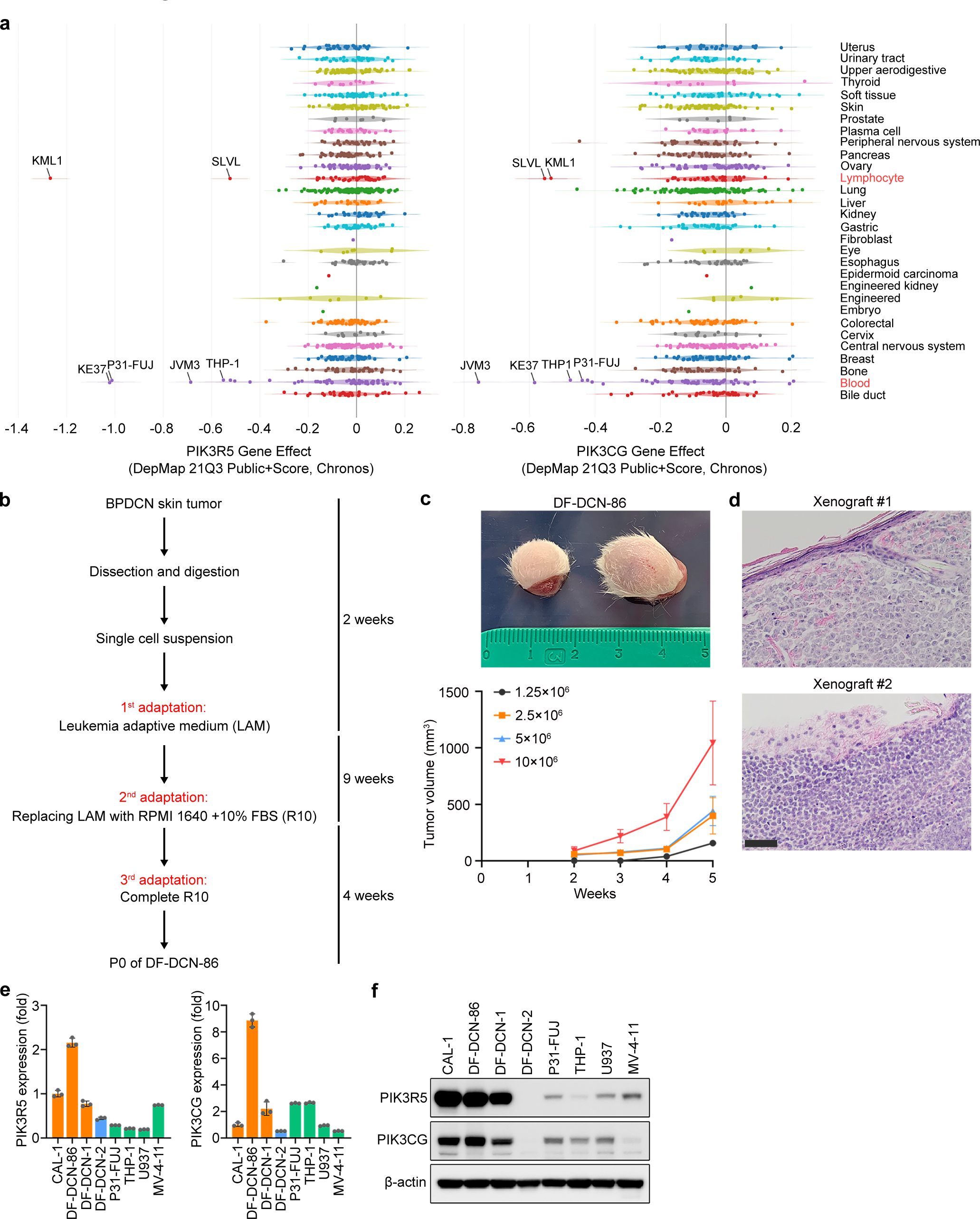
Newly established BPDCN cell lines maintain features of patient samples. **a,** DepMap gene effects of *PIK3R5* and *PIK3CG* in all cancer types. **b,** Flowchart showing the strategy of using LAM to establish new BPDCN cell lines. **c,** Representative images and tumor volume of the intradermal tumors formed by DF-DCN-86 in NSG mice. Data are means ± S.E.M. from 2 biological replicates. **d,** Hematoxylin and eosin staining of the tissue sections from intradermal tumors formed by DF-DCN-86. Scale bar, 50 μm. **e,** RT-qPCR of PIK3R5 and PIK3CG mRNA in leukemia cell lines. Data are means ± S.D. from 3 technical replicates. **f,** Western blotting for PIK3R5 and PIK3CG in leukemia cell lines.

**Extended Data Figure 3.**
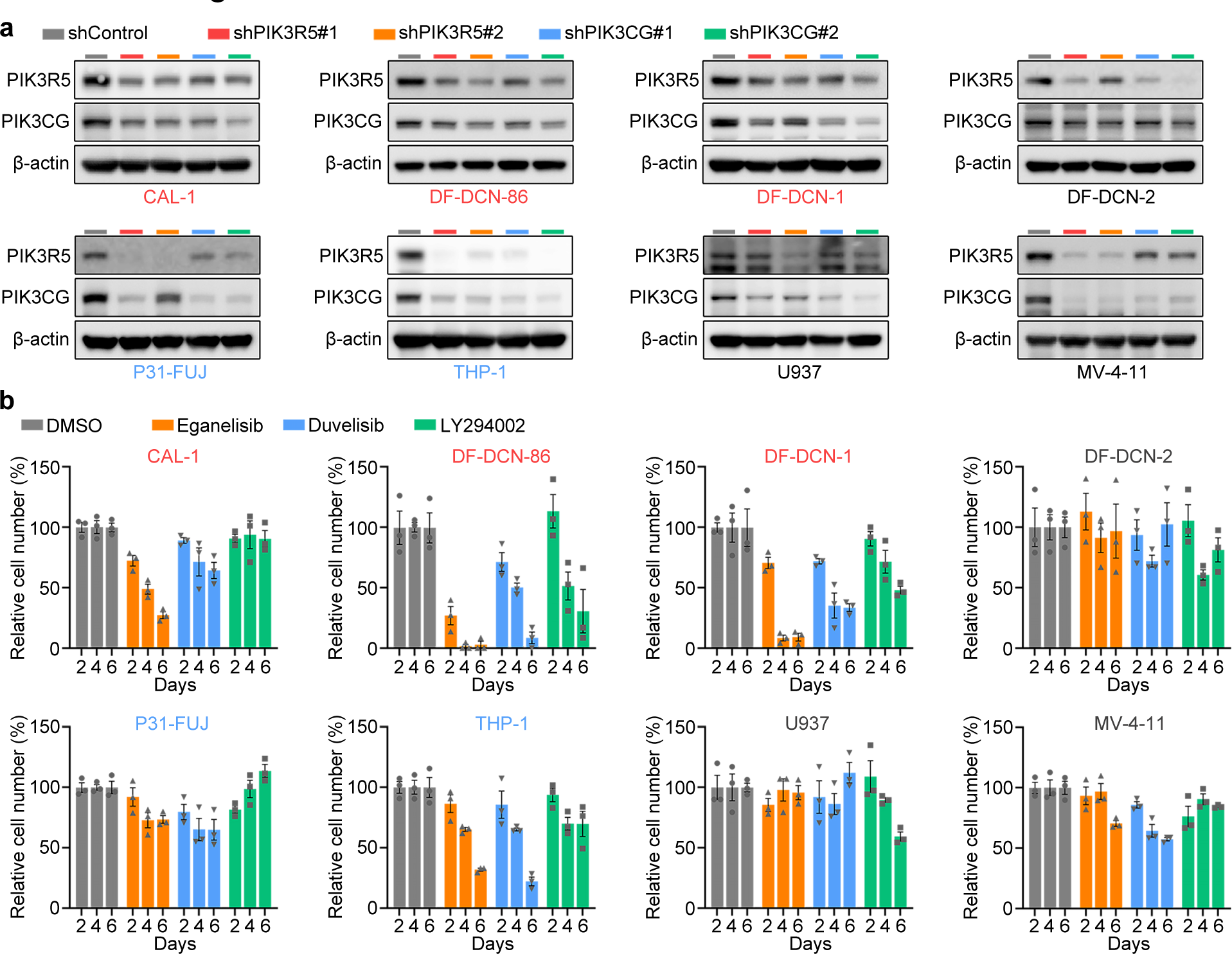
Most BPDCN cell lines are sensitive to PI3Kγ inhibition. **a,** Western blotting for PIK3R5 and PIK3CG in various leukemia cell lines after depletion of PIK3R5 or PIK3CG. **b,** Relative cell numbers of various leukemia cell lines after 2, 4, and 6 days of treatment with eganelisib, duvelisib, or LY294002 (each at 1 μM) as compared to control cells. Strongly sensitive cell lines are labeled in red, moderately sensitive cell lines are labeled in blue, and insensitive cell lines are labeled in grey. Data are means ± S.E.M. from 3 biological replicates.

**Extended Data Figure 4.**
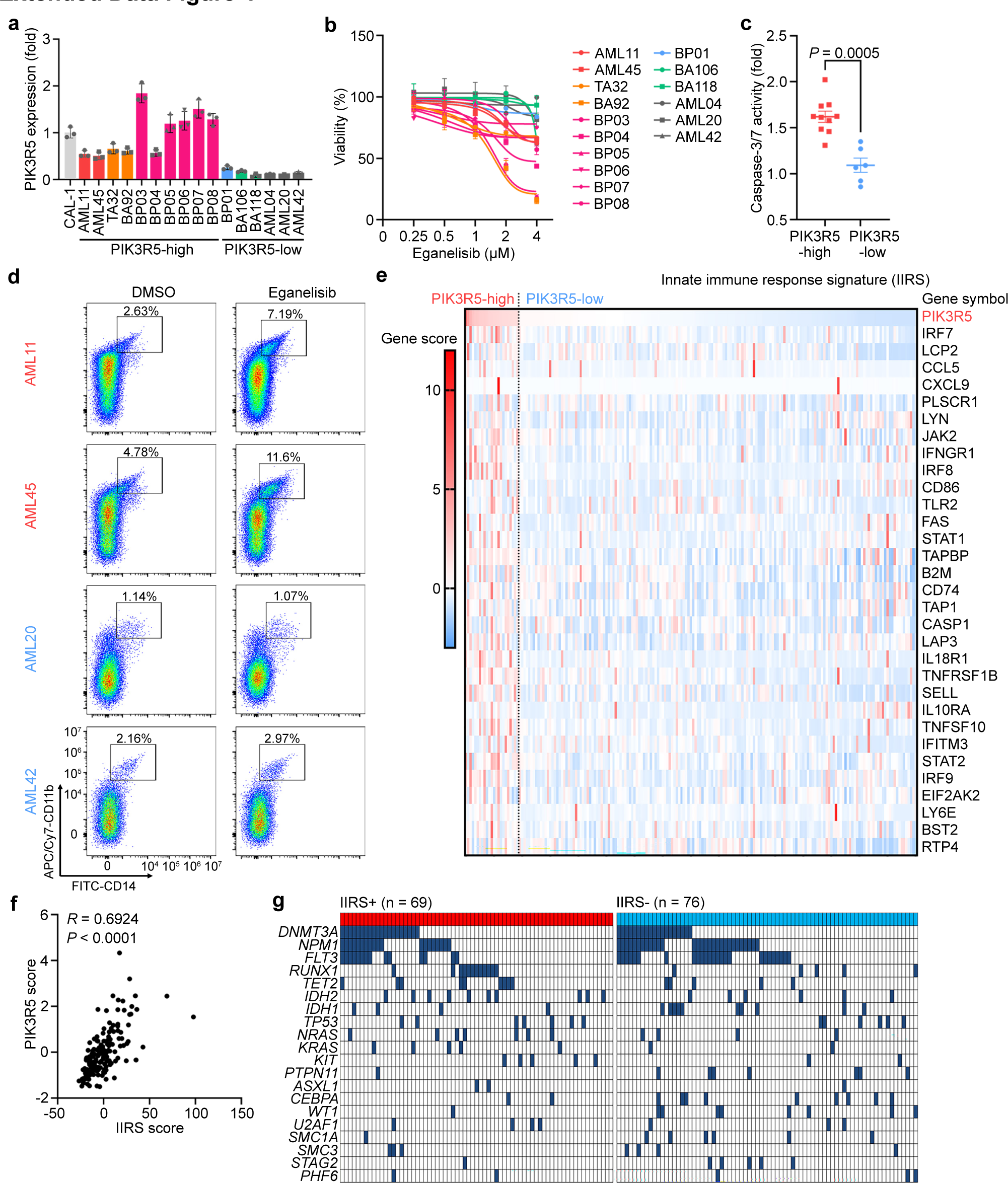
Eganelisib selectively induces apoptosis and promotes differentiation of leukemias with elevated PIK3R5. **a,** RT-qPCR of PIK3R5 mRNA in leukemia PDXs. Data are means ± S.D. from 3 technical replicates. **b,** Inhibition curve of leukemia PDXs after ex vivo treatment with increasing doses of eganelisib (0-4 μM) for 72 hours. Data are means ± S.D. from 3 technical replicates. **c,** Relative caspase-3/7 activities in leukemia PDXs after ex vivo treatment with eganelisib (1 μM) for 72 hours as compared to vehicle control (DMSO). Data are means ± S.E.M. with Mann-Whitney test. **d,** Flow cytometry of CD11b and CD14 in leukemia PDXs after ex vivo treatment with eganelisib (1 μM) or vehicle control (DMSO) for 72 hours. **e,** Heatmap showing expression scores of IIRS genes. The IIRS score was determined by first transforming the relative expression of each gene to values resulting in a mean of 0 and SD of 1 across all patients, then summing up individual gene scores to obtain the overall score. **f,** Correlation analysis of PIK3R5 and IIRS expression scores in leukemia PDX samples. Spearman correlation coefficients are shown. **g,** Comutation plot showing gene mutations of TCGA AML cases with positive or negative IIRS scores.

**Extended Data Figure 5.**
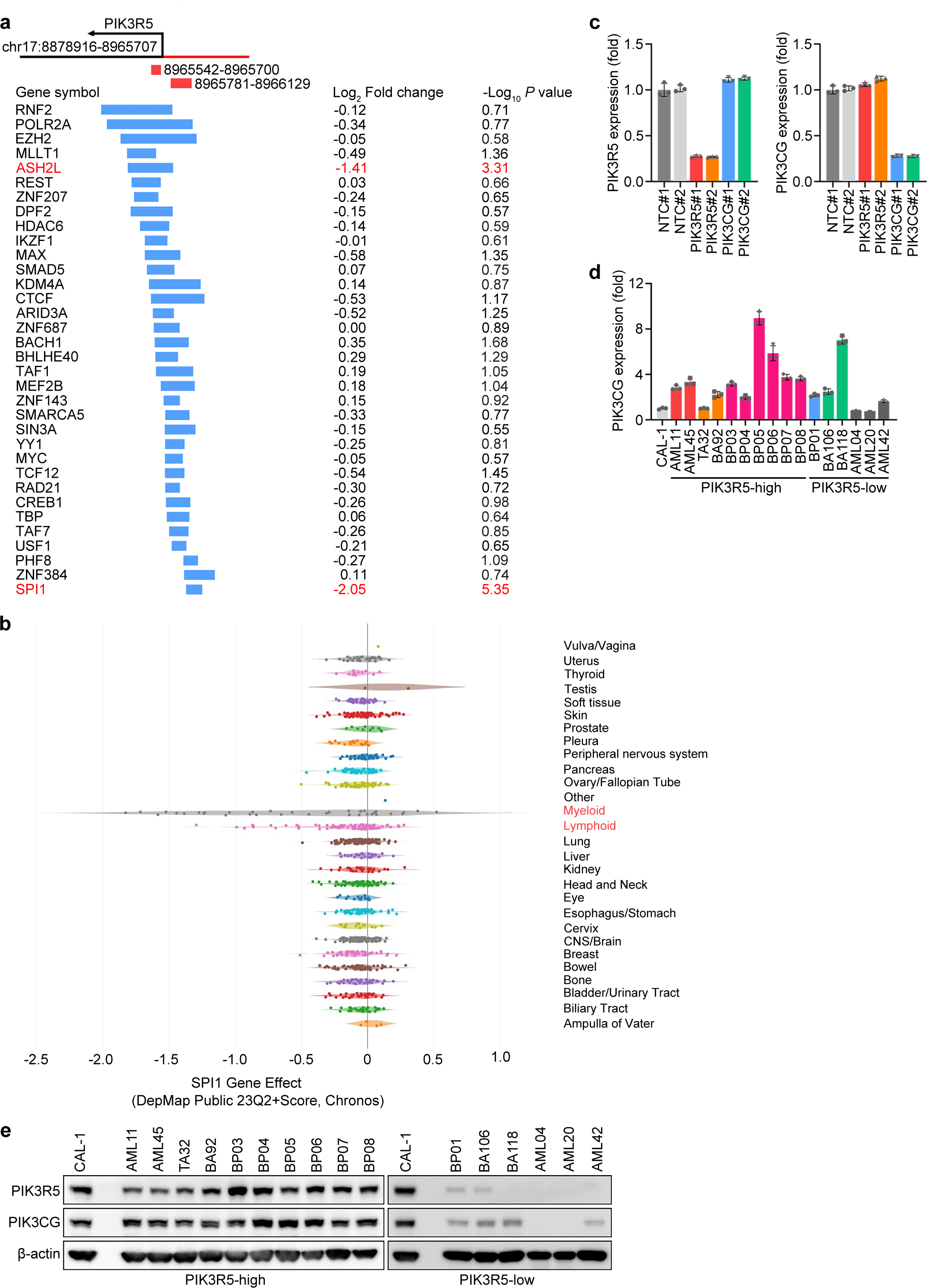
PU.1 activates *PIK3R5* transcription. **a,** Schematic showing the overlapping areas between *PIK3R5* promoter regions and transcriptional regulators from ChIP-seq data in the ENCODE database. Each gene’s log_2_ fold change and -log_10_ *P* value from the genome-wide CRISPRi screen (Fig. 1a; day 21 vs day 0) are shown on the right side. **b,** DepMap gene effect of *SPI1* targeting across cancer types. **c,** RT-qPCR of PIK3R5 and PIK3CG mRNA in CAL-1 cells expressing nontargeting control (NTC#1 and NTC#2) or PIK3R5/PIK3CG targeting CRISPRi guide RNAs (PIK3R5#1, PIK3R5#2, PIK3CG#1, and PIK3CG#2). Data are means ± S.D. from 3 technical replicates. **d,** RT-qPCR of PIK3CG mRNA in leukemia PDXs. Data are means ± S.D. from 3 technical replicates. **e,** Western blotting for PIK3R5 and PIK3CG in various leukemia PDX samples.

**Extended Data Figure 6.**
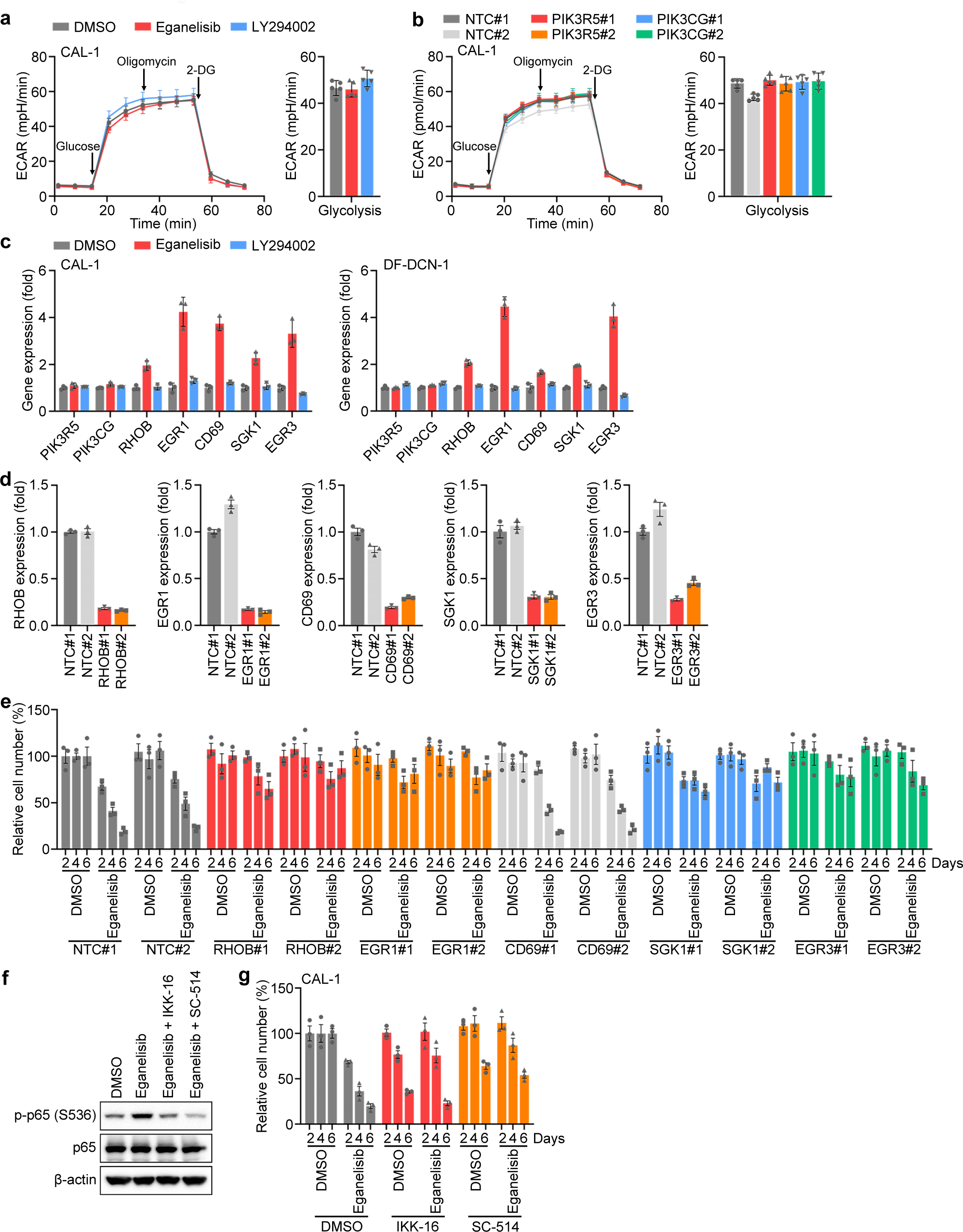
PI3Kγ inhibition activates an NFκB-mediated transcriptional network of tumor suppressor genes. **a,** Seahorse assays measuring glycolysis in the indicated leukemia cell lines upon treatment with eganelisib or LY294002 (each at 1 μM for 48 hours). Data are means ± S.D. from 5 technical replicates. **b,** Seahorse assays measuring glycolysis in control, PIK3R5 depleted, or PIK3CG depleted CAL-1 cells. Data are means ± S.D. from 5 technical replicates. **c,** RT-qPCR of gene expression in CAL-1 cells upon treatment with eganelisib or LY294002 (each at 1 μM for 48 hours). Data are means ± S.D. from 3 technical replicates. **d,** RT-qPCR of gene expression in CAL-1 cells expressing nontargeting control or gene-targeting CRISPRi guide RNAs. Data are means ± S.D. from 3 technical replicates. **e,** Relative numbers of CAL-1 cells expressing nontargeting control or gene-targeting CRISPRi guide RNAs. Data are means ± S.E.M. from 3 biological replicates. **f,** Western blotting for p65 phosphorylation in CAL-1 cells treated with eganelisib (1 μM), together with or without NFkB inhibitors IKK-16 (0.5 μM) or SC-514 (20 μM), for 2 days. **g,** Relative numbers of CAL-1 cells after treatment with eganelisib (1 μM), with or without NFkB inhibitors IKK-16 (0.5 μM) or SC-514 (20 μM), for 2, 4, and 6 days. Data are means ± S.E.M. from 3 biological replicates.

**Extended Data Figure 7.**
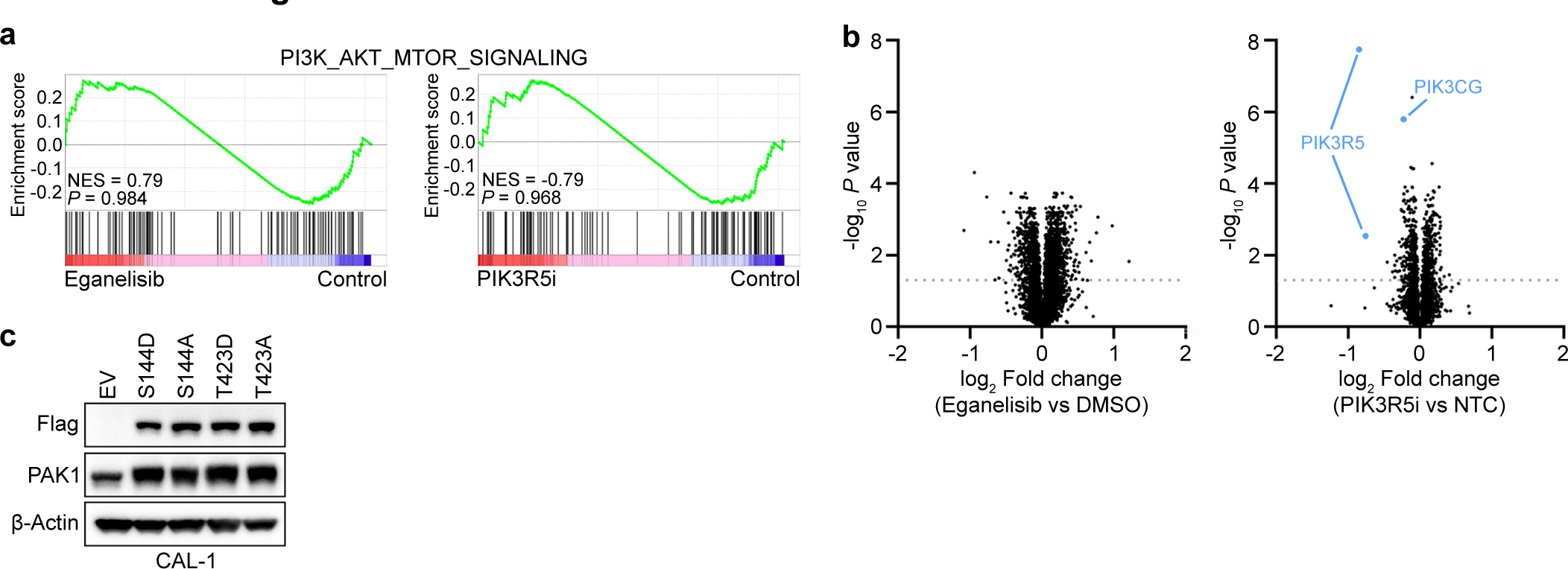
PI3Kγ inhibition does not affect the PI3K-AKT-mTOR signaling pathway. **a,** GSEA indicating that the PI3K-AKT-mTOR signaling pathway downstream genes were not significantly affected upon eganelisib treatment (1 μM for 48 hours) or PIK3R5 depletion (PIK3R5i) in CAL-1 cells. **b,** Volcano plot showing the upregulated or downregulated proteins by mass spectrometry in CAL-1 cells treated with eganelisib or depleted of PIK3R5 (PIK3R5i). PIK3R5 (two dots represent different PIK3R5 Uniprot protein IDs Q8WYR1 and J3KSW1, respectively) occurred as the most significantly downregulated protein, confirming the knockdown efficiency of PIK3R5. PIK3CG protein level was also moderately decreased due to its reduced stability after PIK3R5 depletion. Adjusted *P* values are shown. **c,** Western blotting for Flag and PAK1 in CAL-1 cells expressing empty vector (EV) or Flag-tagged PAK1 mutants.

**Extended Data Figure 8.**
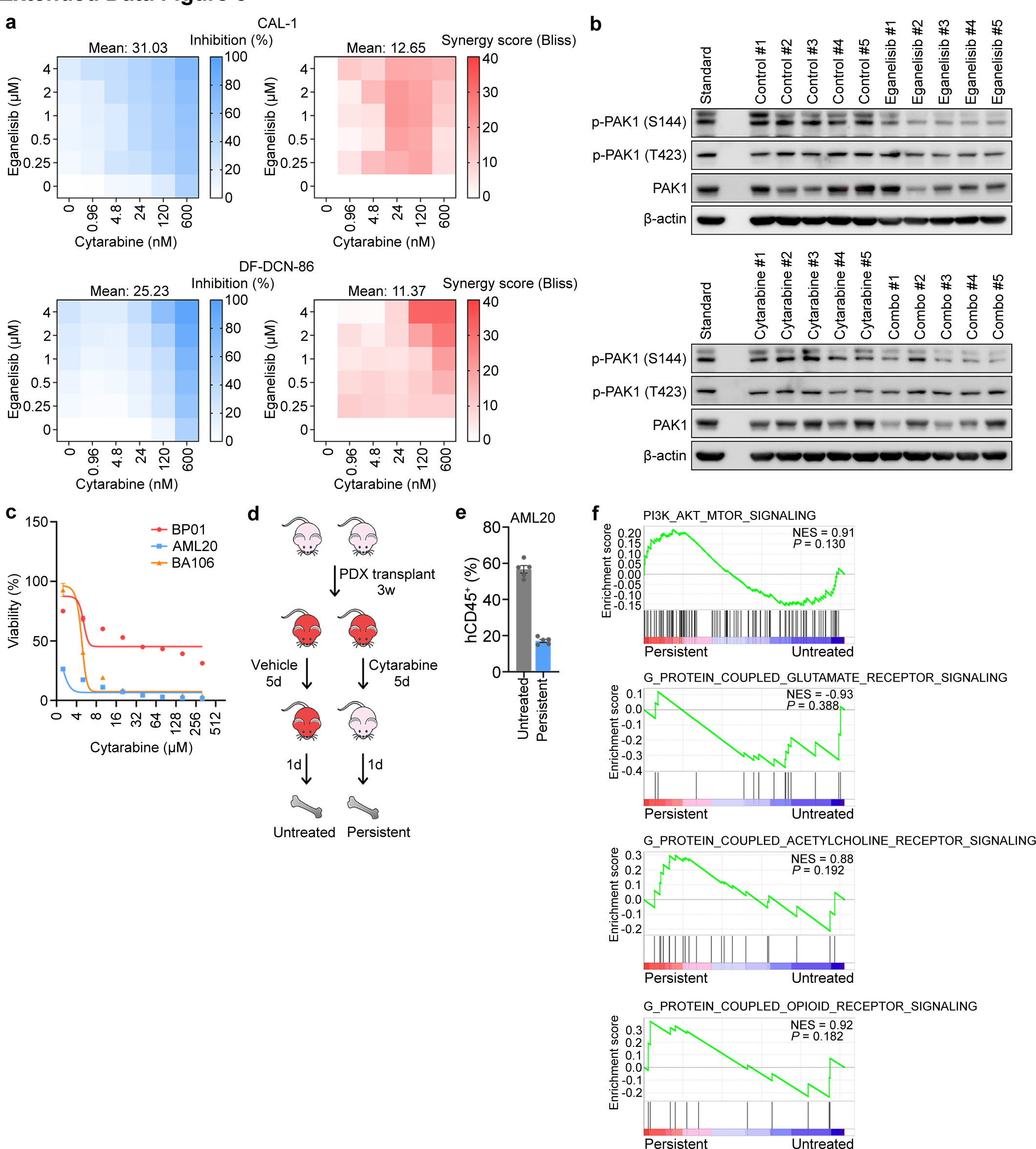
Eganelisib synergizes with cytarabine in leukemia treatment in vivo. **a,** Heatmaps showing inhibition percentages and synergy scores of combined eganelisib and cytarabine treatment in CAL-1 and DF-DCN-86 cells. **b,** Western blotting for PAK1 phosphorylation in DF-DCN-86 intradermal xenografts. The standard sample was created by mixing an equal amount from each of the 5 control samples. **c,** Inhibition curve of leukemia PDXs after ex vivo treatment with increasing doses of cytarabine (0-320 μM) for 72 hours. Data are means ± S.D. from 3 technical replicates. **d,** Schematic showing the strategy for establishing a cytarabine-persistent PDX model using AML20. **e,** Flow cytometry for human CD45^+^ cell percentage in bone marrow samples from untreated or cytarabine-treated mice. Data are means ± S.E.M. from 5 biological replicates. **f,** GSEA indicating that neither the PI3K-AKT-mTOR signaling pathway nor other non-purinergic G protein-coupled receptor signaling pathways were significantly elevated in cytarabine-persistent AML20 cells.

